# The m^6^A Reader ECT1 Mediates Seed Germination via the DAG2-ECT1-PHYB Regulatory Cascade in *Arabidopsis*

**DOI:** 10.1101/2024.07.24.604550

**Authors:** Zenglin Li, Yuhang Ma, Wen Sun, Pengjun Ding, Yifan Bu, Yuhong Qi, Chengchao Jia, Beilei Lei, Chuang Ma

## Abstract

The *N*^6^-methyladenosine (m^6^A) RNA modification, a significant epitranscriptomic mark, is integral to plant growth and development. m^6^A reader proteins are capable of recognizing m^6^A modifications to impact RNA metabolic and biological processes. Here, we discovered ECT1 as an m^6^A reader protein that directly binds to the m^6^A site and plays a role in positively regulating seed germination in the context of gibberellins (GAs). ECT1 undergoes m^6^A modification to ensure its own stability and engages in self-interaction to enhance its protein stability. Moreover, ECT1 establishes a regulatory circuit with DAG2, which has been reported to play a positive regulatory role in GA-mediated seed germination. We found that DAG2 directly binds to the *ECT1* promoter to control its transcription. ECT1 modulates the mRNA stability of *DAG2*. Furthermore, we identified *PHYB*, which encodes a key photoreceptor controlling GA-regulated seed germination, as a common downstream target of DAG2 and ECT1. DAG2 and ECT1 regulate the expression of *PHYB* at the transcriptional and post-transcriptional levels, respectively. ECT1 binds directly to *PHYB*, thereby influencing its stability. While DAG2 binds to the *PHYB* promoter to regulate its transcription. Together, these results unveil that ECT1 participates in the m^6^A signaling pathway through complex and multifaceted molecular mechanisms, broadening the spectrum of m^6^A reader functions and deepening our insight into the m^6^A signaling network in *Arabidopsis*.

## Introduction

m^6^A modification is among the most abundant and important RNA chemical modifications, playing a crucial role in various biological functions (Shao et al., 2021). The m^6^A modification in eukaryotic mRNA is a dynamic process involving the installation of methyl groups by RNA methyltransferases referred to as "writers" and the removal of these methyl groups by demethylases referred to as "erasers". Furthermore, m^6^A residues are acknowledged and interpreted by RNA-binding proteins (RBPs) referred to as "readers" (Jia et al., 2011; Zheng et al., 2013; Meyer and Jaffrey, 2014; Hu et al., 2019; Shen et al., 2019; Arribas-Hernández and Brodersen, 2020; Zhang et al., 2022a; Amara et al., 2023). The m^6^A modification has been observed to be highly concentrated in the 3’-UTR and near stop codons of eukaryotic mRNAs, indicating its potential involvement in post-transcriptional regulation of gene expression (Meyer et al., 2012; Liu et al., 2020; Guo et al., 2022; Zhou et al., 2022; Miao et al., 2022). While the presence of m^6^A has been known since the 1970s in mammals, prokaryotes, and viruses, its precise function remained largely elusive until recent advancements in identifying its writers, erasers, and readers and developing high throughput m^6^A sequencing technologies (Dubin et al., 1975; Wei et al., 1975; Perry et al., 1975; Meyer et al., 2012; Xu et al., 2015).

Currently, m^6^A has been revealed to play functional roles in modulating various aspects of mRNA metabolism including pre-mRNA splicing, mRNA stability, translation efficiency, alternative polyadenylation, alternative splicing, nuclear-to-cytoplasmic mRNA export, and microRNA processing (Wang et al., 2014; Huang et al., 2018; Wang et al., 2015; Roundtree et al., 2017; Zhou et al., 2022). The importance of m^6^A modifications in gene regulation and biological functions has become increasingly evident (Sharma et al., 2023; Shen et al., 2023). m^6^A modification has been found to modulate shoot stem cell proliferation, trichome branching, floral transition, abscisic acid (ABA) response, chilling tolerance, drought and salt tolerance, photosynthesis, and nitrate signaling in *Arabidopsis* (Wang et al., 2021; Wang et al., 2022a; Jiang et al., 2023; Wei et al., 2018; Shen et al., 2016; Wang et al., 2022b; Zhou et al., 2022; Zhang et al., 2022b; Hou et al., 2021). As well as rice sporogenesis and grain yield, the development of trichomes and roots in poplar, and fruit ripening of strawberry (Zhang et al., 2019; Yu et al., 2021; Lu et al., 2020; Zhou et al., 2021).

YT521-B homology (YTH) domain-containing proteins, found across various eukaryotes, exhibit a conserved YTH domain with an aromatic cage structure. This specialized structure enables them to selectively recognize and bind RNA molecules containing m^6^A modification, making them the principal "readers" of m^6^A (Zhang et al., 2010; Luo and Tong, 2014; Theler et al., 2014; Xu et al., 2014; Patil et al., 2018; Amara et al., 2024; Song et al., 2023). In *Arabidopsis*, thirteen YTH family proteins have been computationally identified, among which ECT2, ECT3, ECT4, ECT8, ECT12, and CPSF30-L, have been characterized as m^6^A reader proteins (Wei et al., 2018; Song et al., 2023; Amara *et al*., 2024; Hou et al., 2021; Cai et al., 2024). CPSF30-L is a nuclear m^6^A reader protein that regulates alternative polyadenylation (APA) to control nitrate signaling; ECT2, ECT3, and ECT4 redundantly function in seed germination, leaf growth and organogenesis; ECT12 affects abiotic stress responses by modulating mRNA stability (Hou et al., 2021; Arribas-Hernández et al., 2020; Song et al., 2023; Amara et al., 2024; Cai et al., 2024).

Over the last decade, there has been a significant surge in studies focused on characterizing and comprehending m^6^A modifications in the field of plant biology. Despite the extensive documentation of m^6^A’s biological functions in essential plant processes, the precise molecular mechanism governing its regulatory roles still lacks comprehensive understanding (Zhou et al., 2022; Shen et al., 2023; Song et al., 2023; Wang et al., 2024). Recently, Lee et al. reported that ECT1 can function as a reader protein and play a significant role in SA-mediated stress responses. This revelation has unveiled further possibilities for ECT1 as a reader protein (Lee et al., 2024).

In this study, we employed *in vivo* and *in vitro* experiments to validate that ECT1 can directly bind to m^6^A-modified RNAs, thereby exerting its reader protein function, confirming ECT1 as an m^6^A reader protein. We have identified ECT1 as an essential m^6^A reader protein, which positively regulates seed germination. ECT1 exhibits multiple roles within the complex m^6^A signaling pathway, operating through various intricate molecular mechanisms. ECT1 undergoes m^6^A modification to ensure its own stability and that of *DAG2* and *PHYB*, thereby ensuring the proper functionality of the m^6^A signaling pathway. Furthermore, ECT1 participates in self-interaction, a mechanism that reinforces its protein stability. ECT1 forms a regulatory loop with DAG2, where DAG2 directly interacts with the *ECT1* promoter to govern its transcription; ECT1, in turn, affects the mRNA stability of *DAG2* via m^6^A modifications. Moreover, we have identified *PHYB* as a shared downstream target of both DAG2 and ECT1. ECT1 and DAG2 regulate *PHYB* via an independent pathway.

## Results

### ECT1 is an m^6^A reader protein

Since the direct interaction between ECT1 and m^6^A has not been previously investigated, we conducted an Electrophoretic Mobility Shift Assay (EMSA) to examine whether ECT1 can bind to m^6^A-modified RNAs. Initially, we expressed the recombinant GST-ECT1 protein in *E. coli* and purified it. We synthesized RNAs labeled with 5C-biotin, which was reported to be bound by m^6^A reader contained the *UGUAA* motif known to be bound by m^6^A reader proteins (Wei et al., 2018), for use in the EMSA. The EMSA results revealed that ECT1 caused a distinct shift in the migration pattern of the m^6^A-modified synthetic RNA, indicating its ability to bind to m^6^A-modified RNAs (Fig. 1a). Conversely, no change was observed when non-m^6^A-modified synthetic RNA was utilized (Fig. 1a). To serve as a negative control, the GST tag trigger factor was employed, and it did not exhibit any binding to the RNAs as expected (Fig. 1a). These findings suggest that ECT1 has the potential to function as a reader protein for m^6^A modifications.

**Fig. 1.**
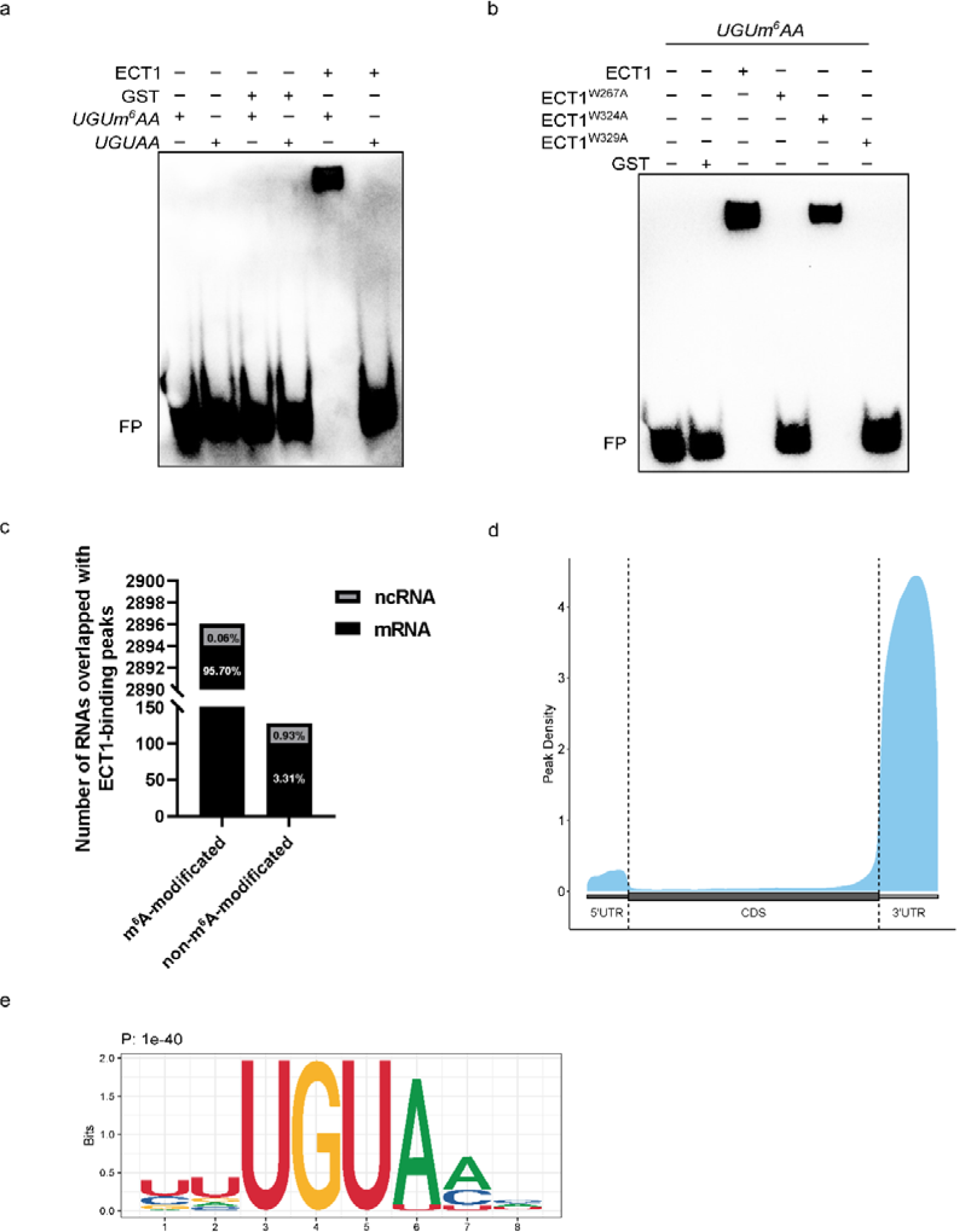
RNA-binding ability of ECT1. **(a)** EMSA assay. 5_-Biotin-labeled synthetic RNA with or without m^6^A modification motif *UGUAA* were incubated with GST-ECT1 protein or GST protein alone (negative control). Samples were analysed by native PAGE. Gels were blotted onto nylon membranes and signals were detected by streptavidin-coupled horseradish peroxidase and ECL. FP, free probe. **(b)** EMSA assay. Biotin-labeled m^6^A-modified (*UGUm^6^AA*) RNA fragment was incubated with GST-ECT1 and different mutants of GST-ECT1 (amino acid W-267 mutated to A, GST-ECT1^W267A^; amino acid W-324 mutated to A, GST-ECT1^W324A^; amino acid W-329 mutated to A, GST-ECT1^W329A^) or GST (negative control). Samples were analysed by native PAGE. Gels were blotted onto nylon membranes and signals were detected by streptavidin-coupled horseradish peroxidase and ECL. FP, free probe. **(c)** Number of overlaps with the ECT1 binding peak for each type of RNA. **(d)** Distribution of ECT1-binding peaks along with the 5’UTR, CDS, and 3’UTR regions of protein-coding genes. **(e)** Motif identified by HOMER based on ECT1-binding peaks located within protein-coding genes.

Previous studies have highlighted the importance of the aromatic cage within the YTH domain in forming a hydrophobic binding pocket, enabling it to recognize methyl groups of m^6^A residues (Wei et al., 2018; Luo et al., 2023). Among the amino acid residues of ECT1, Trp267, Trp324, and Trp329 are particularly important in establishing this aromatic cage (Supplementary Fig. 1). Based on this, we hypothesize that these three tryptophan residues of ECT1 directly participate in the binding of ECT1 to m^6^A sites. To investigate this, we designed a mutated form of ECT1, named ECT1^W267A^, ECT1^W324A^, and ECT1^W329A^, where the tryptophan residues at positions 267, 324, and 329 were replaced with alanine, respectively. Recombinant GST-ECT1^W267A^, GST-ECT1^W324A^, and GST-ECT1^W329A^ proteins were expressed and purified from *E. coli*. We then conducted EMSA using synthesized RNAs labeled with 5C-biotin, containing either m^6^A or A. The EMSA analysis revealed that ECT1^W267A^ and ECT1^W329A^ almost completely abolished the binding function (Fig. 1b). At the same time, ECT1^W324A^ exhibited a decreased m^6^A binding capacity compared to the wild-type ECT1 protein (Fig. 1b). These results collectively confirm that ECT1 binds to RNA transcripts containing m^6^A sites, solidifying its role as an m^6^A reader protein.

To confirm the recognition of m^6^A sites by ECT1 in plants, we generated transgenic *Arabidopsis* plants (*35S: ECT1-FLAG*) expressing an ECT1-FLAG fusion protein under the control of the CaMV 35S promoter in the wild-type (Col-0) background (Supplementary Fig. 2). Subsequently, we performed an RNA immunoprecipitation-seq (RIP-seq) experiment using the ECT1-FLAG plants (ECT1OE-8). From the RIP-seq analysis, the majority of transcripts targeted by ECT1 were found to be mRNA molecules, with a small fraction (0.99%) being noncoding RNA (Fig. 1c). Furthermore, when we compared the RIP-identified ECT1 targeted sites with the confidence m^6^A methylation peaks from wild-type seedings from IPOP database (Miao et al., 2022; Huang et al., 2023), we observed that 95.76% (2896 out of 3024) of the ECT1-targeted peaks were modified with m^6^A (Fig. 1c). These findings provide evidence for ECT1’s recognition of m^6^A on RNA transcripts in plants. We also explored additional characteristics of the ECT1-targeted transcripts and found that many of the ECT1-targeted sites were located within the 3’ UTR of RNA transcripts (Fig. 1d). By utilizing HOMER (Hypergeometric Optimization of Motif Enrichment) to perform motif enrichment analysis of ECT1 binding peaks, we identified a highly enriched motif: UGUAH (H=A>C>U), (Fig. 1e), where UGUAY (Y=C>U) is a plant-specific m^6^A motif (Wei et al., 2018). These findings firmly support the notion that ECT1 acts as a m^6^A reader protein in *Arabidopsis*.

### ECT1 binds to its own mRNA for the stability of its mRNA

Among the transcripts targeted by ECT1, we found that *ECT1* mRNA itself is also a target of ECT1 (Supplementary Tab. 5), and capable of being modified by m^6^A from our comparison of the RIP-identified ECT1 target sites with the confident m^6^A methylation peaks from wild-type seedlings in the IPOP database (Miao et al., 2022; Huang et al., 2023). To further investigate the interaction between ECT1 and its mRNA, an RIP-qPCR assay was performed using 4-day-old ECT1-FLAG overexpressing plants and wild-type plants as controls. The RIP-qPCR analysis revealed a significant enrichment of *ECT1* in the ECT1-FLAG transgenic plants compared to the wild-type (Fig. 2a). This finding confirmed the ability of ECT1 to bind to the m^6^A-modified *ECT1* transcripts in *Arabidopsis*.

**Fig. 2.**
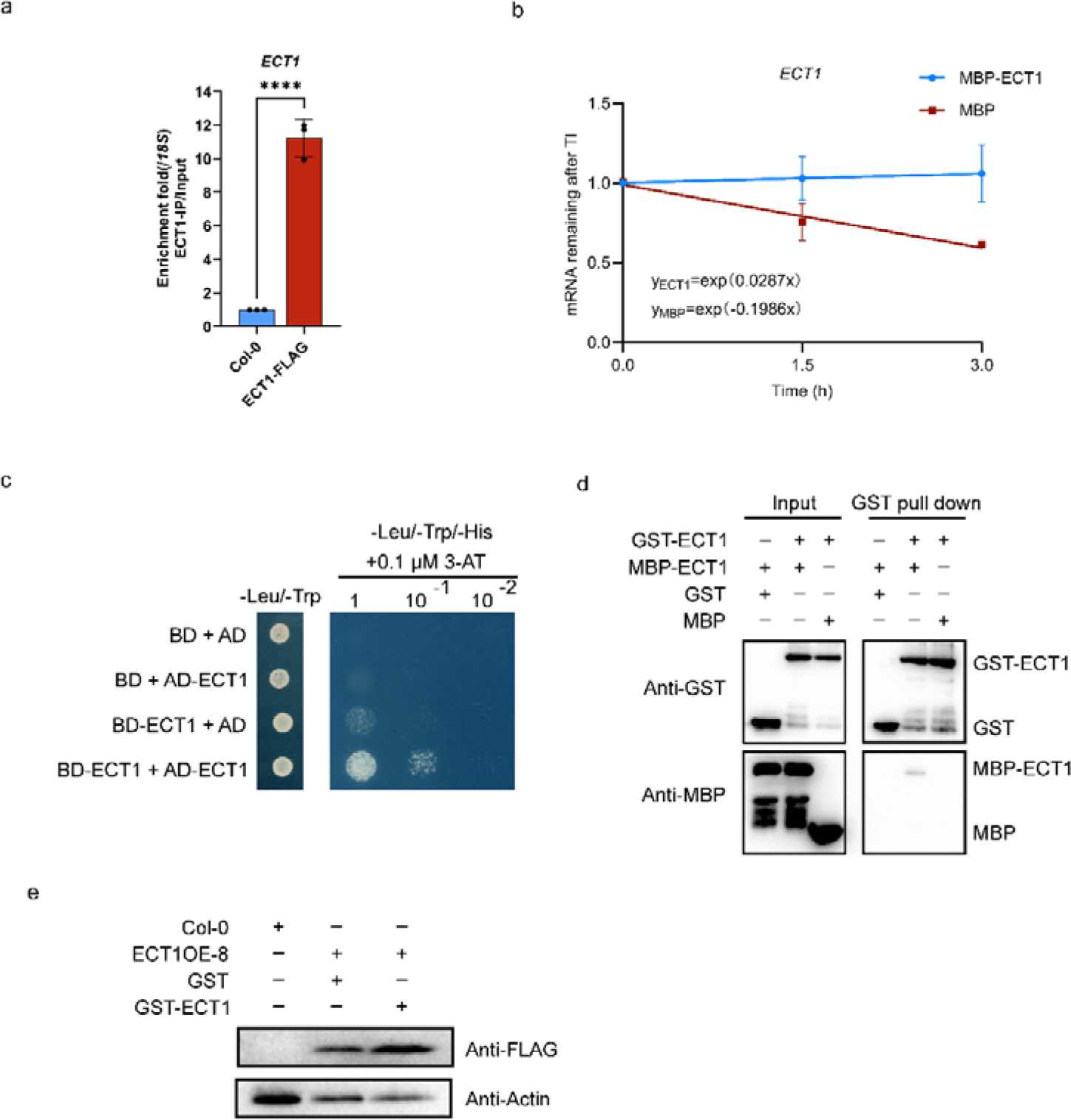
ECT1 promotes the stability of its own mRNA and protein. **(a)** RIP-qPCR assay. 4-day-old seedings of Col-0 *p35S: ECT1-FLAG* (line 8) and wild type Col-0 (negative control) were used in RIP assay. Chromatin was isolated and ECT1-FLAG bound RNA fragments were purified using anti-FLAG coupled magnetic beads. *ECT1* RNA fragments in the input and eluate fractions were detected using qPCR and specific primers. *18S* RNA was used for normalization. Values show enrichment of *ECT1* RNA fragments in the eluate fraction compared to the input fraction and are means of three replicates ± SD. Significant differences as determined by student’s t-test, ****: *p*<0.0001. **(b)** mRNA stability assay. mRNA lifetimes of *ECT1* in RNA from 4-day-old wild-type seedlings incubated with MBP-ECT1 or MBP protein and treated with actinomycin D. TI, transcription inhibition. *18S* RNA was used for normalization. Data are means ± SD for three replicates. **(c)** Y2H protein-protein interaction assay. AD-ECT and BD-ECT1 were expressed in yeast. Yeast cells were diluted 10 and 100-fold and then cultured on CSM-L-T plates or CSM-L-T-H supplemented with 0.1 μM 3-AT plates. **(d)** GST Pull-down assay. Purified MBP-ECT1 or MBP was incubated with GST-ECT1 or GST proteins. GST-ECT1 bound proteins were purified using anti-GST coupled magnetic beads, followed by immunoblot analysis using GST and MBP antibodies. **(e)** Protein stability assay. GST-ECT1 and GST protein were co-incubated with protein extracts from 4-day-old Col-0 *p35S: ECT1-FLAG* seedlings for two hours, followed by analysis using SDS-PAGE and immunoblotting with anti-FLAG. Anti-Actin was used to detect ACTIN as a loading control. Col-0 was included as a negative control.

Given the evidence that mRNA m^6^A modification can influence the stability of individual mRNAs in *Arabidopsis* (Wei et al., 2018; Tang et al., 2022), we conducted transcription inhibition (TI) assays to examine whether ECT1 is involved in regulating the stability of *ECT1* transcript. We assessed the durations of *ECT1* transcripts by inhibiting transcription using actinomycin D. Simultaneously, we introduced purified recombinant proteins MBP-ECT1 and MBP from *E. coli* in this experiment. Through this assay, we determined that the presence of MBP-ECT1 led to increased stability of *ECT1* transcript compared to the control with MBP tag alone (Fig. 2b). Consequently, our findings collectively demonstrate the crucial role of ECT1 in stabilizing *ECT1* transcripts.

### The ECT1 protein forms self-associations to preserve its stability

Previously, there was no report of any self-interaction among the 13 YTH proteins in *Arabidopsis*. To investigate the possibility of self-interaction of ECT1, we conducted two experiments. Initially, a yeast two-hybrid assay was performed, where yeast strains co-transformed with AD-ECT1 and BD-ECT1 were able to grow on a selective medium (Fig. 2c), indicating the occurrence of protein-protein interactions. To further validate this finding, *in vitro* GST pull-down assays were carried out using purified recombinant proteins GST-ECT1 and MBP-ECT1 from *E coli*. The results showed that MBP-tagged ECT1 interacted with GST-tagged ECT1 but not with GST alone (Fig. 2f). Similarly, GST-tagged ECT1 interacted with MBP-tagged ECT1 but not with MBP alone (Fig. 2d). This suggests that ECT1 can form an ECT1-ECT1 complex through direct protein-protein interactions.

Furthermore, an intriguing observation was made during our study, where we noticed that the presence of ECT1-GST led to an increase in ECT1 protein levels in comparison to GST alone (Fig. 2e). This suggests that apart from promoting *ECT1* mRNA stability, ECT1 also enhances the stability of ECT1 protein.

### ECT1 is a positive regulator of GA-triggered germination

To investigate the biological functions of ECT1, we obtained the *ect1-1* mutant (SALK_073479C) (Supplementary Fig. 3) and screened for its phenotypes. We found that under normal conditions, the wild-type, *ect1-1* mutant showed higher and consistent seed germination rates (Fig. 3a, b). However, in the experiment where seeds were treated with GA_3_, we observed that under 50 μM GA_3_ treatment, the *ect1-1* mutant exhibited a lower seed germination rate compared to the wild-type (Fig. 3a, b). Subsequently, we assessed the sensitivity of the wild-type, *ect1-1* mutant to different concentrations of GA_3_ by measuring the germination rate of seeds. Our observations revealed that the positive effect of GA_3_ on germination is hyposensitive in the *ect1-1* mutant compared to the wild-type (Fig. 3c). These phenotypic results suggest that ECT1 plays a positive role in the regulation of seed germination by gibberellins.

**Fig. 3.**
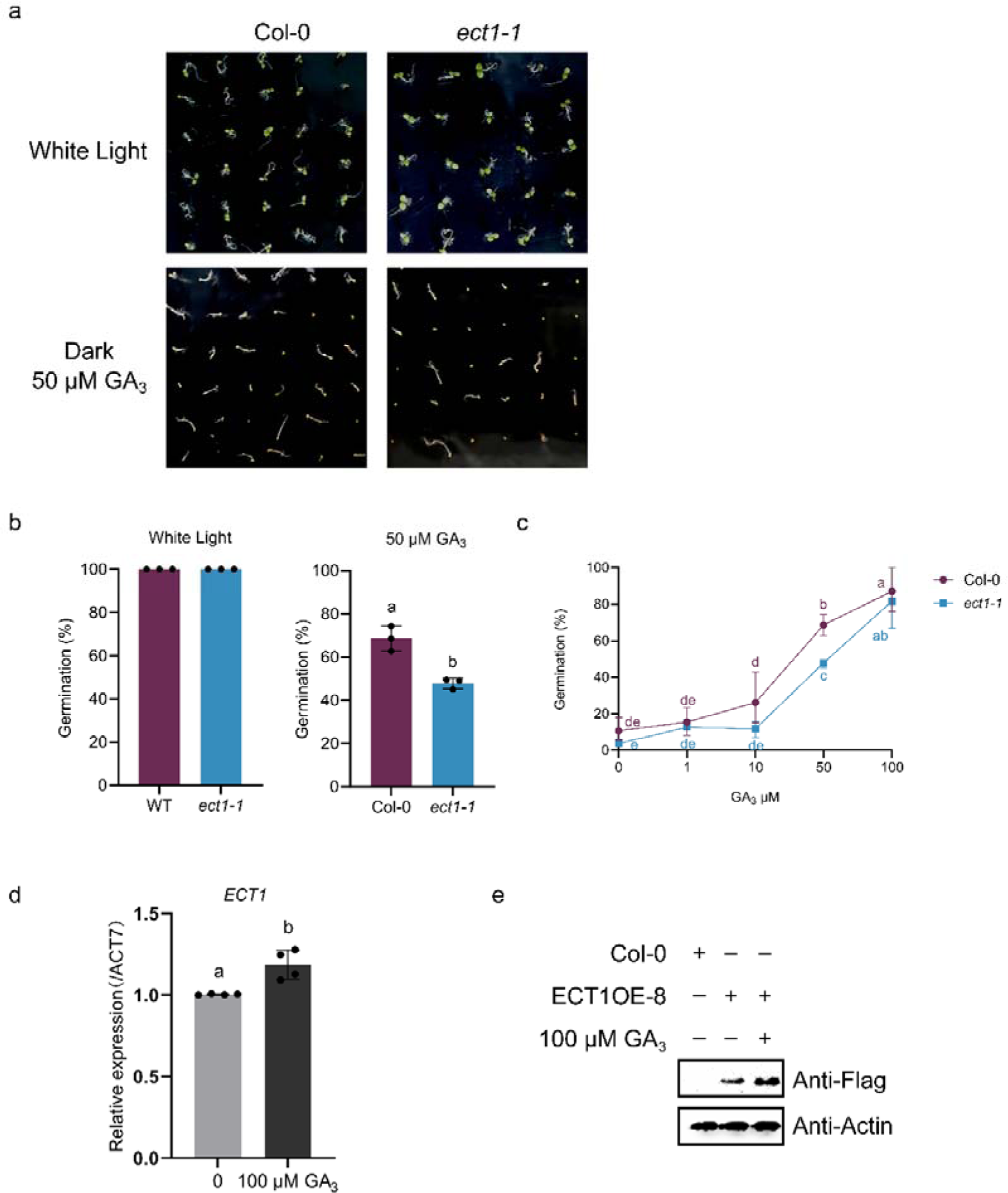
ECT1 is required for seed germination under GA treatment. **(a)** Phenotypic assessment of the GA response in Col-0 and *ect1-1* seeds cultivated on 1/2 MS medium supplemented with 0 or 50 μM GA_3_, under either white light or dark conditions. Photographs representing the growth were captured four days after sowing. **(b)** Analysis of germination percentages in Col-0 and *ect1-1* mutants treated with 0 or 50 μM GA_3_, in either white light or dark conditions. Data from three replicates were averaged. The bar graph represents the average cumulative germination rates from three trials, with error bars indicating ± SD. Significant differences as determined by student’s t-test, **: *p*<0.01. **(c)** The percentage of seed germination for various genotypes in response to GA_3_ treatment. Seeds were sown on MS medium supplemented with varying concentrations of GA_3_, and the germination rates were documented four days post-sowing. The data represent the average of three replicates ± SD. **(d)** RT-qPCR was conducted to quantify *ECT1* mRNA levels in seeds after a 12-hour incubation in the dark with either 0 or 50 μM GA_3_. *Actin7* served as the internal standard. The data represent the average of three replicates ± SD. Significant differences as determined by student’s t-test, **: *p*<0.01. **(e)** Effect of GA_3_ on ECT1-FLAG protein expression in ECT1 overexpression seeds. Seeds from the ECT1OE-8 line were treated with 100 μM GA_3_ for 12 hours in the dark, while control seeds were treated with water. Proteins were then extracted and subjected to SDS-PAGE and immunoblot analysis using an anti-FLAG antibody; anti-Actin was utilized as a loading control to detect ACTIN. Col-0 seeds served as a negative control.

To assess the potential regulatory effects of gibberellins on ECT1, we measured the expression of *ECT1* in seeds germinating on a medium supplemented with 100 uM GA_3_. We found that *ECT1* transcript levels were elevated in seeds germinating in the presence of GA_3_ compared to those without GA_3_ treatment (Fig. 3d). Notably, we observed an increase in ECT1 protein levels in germinating seeds of the ECT1OE-8 when exposed to GA_3_ (Fig. 3e). The ECT1OE-8 constitutively expresses ECT1-FLAG under the control of the CaMV 35S promoter, which is positively regulated by GA_3_ (Fig. 3e). Consequently, GA_3_ not only enhances the expression of *ECT1* but also appears to stabilize the ECT1 protein.

Collectively, these findings indicate that ECT1 contributes positively to the control of seed germination mediated by gibberellins. GA_3_ positively regulates ECT1 at both the transcript level and protein stability, suggesting that gibberellin can play a positive regulatory role in germination by controlling the expression level of ECT1.

### The seed germination regulator DAG2 positively regulate *ECT1*

We complied ten bioinformatics resources to identify potential transcription factors that may regulate *ECT1* (Supplementary Fig. 4a; Supplementary Tab. 6). By intersecting with ECT1-RIP-seq data, we identified four potential transcription factors (e.g., DAG2) that can regulate *ECT1* expression and be regulated by ECT1 (Fig. 4a, Supplementary Fig. 4; Supplementary Tab. 6, 7). Among them, DAG2 is important in positively regulating seed germination (Gualberti et al., 2002; Santopolo et al., 2015). Thus, we expect that ECT1 can form a transcriptional feedback regulatory loop with DAG2 to control seed germination.

**Fig. 4.**
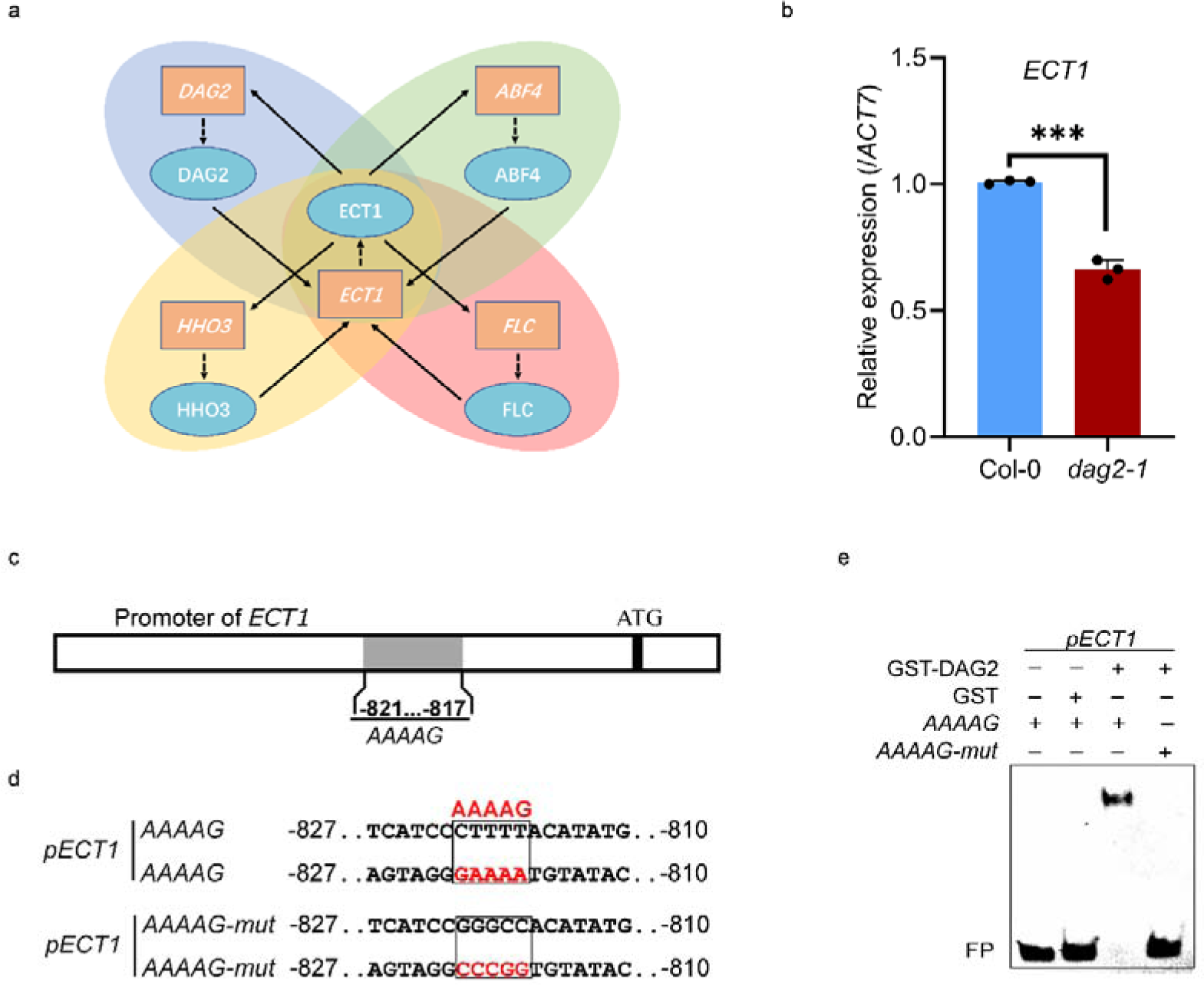
DAG2 induce *ECT1* expression. **(a)** Schematic diagram of ECT1 and transcription factor with potential feedback regulation. Ellipses represent proteins, rectangles represent genes. **(b)** Comparison of ECT1 mRNA expression in 4-day-old Col-0 and *dag2-1* seedlings. *Actin7* served as the reference gene for normalization. The presented data are the average of three replicates, with ± SD shown. Statistical significance was assessed using student’s t-test, with *** indicating *p*<0.001. **(c)** and **(d)** Schematic representation of the *ECT1* promoter. The promoter fragment used for EMSA is shown in (d). **(e)** Biotin-labelled *ECT1* −810…−827 promoter fragments with either a wild-type or mutated *AAAAG* motif were incubated with GST-DAG2, or GST alone (negative control). Samples were analyzed by native PAGE. The gel was blotted onto a nylon membrane and signals were detected by streptavidin-coupled horseradish peroxidase and ECL. FP, free probe.

To support the role of DAG2 in regulating *ECT1* expression, we obtained the *dag2-1* mutant (SALK_201125C) and it exhibits a phenotype similar to that of *ect1-1* (Supplementary Fig. 5). Then we quantified *ECT1* expression in wild-type, and *dag2-1* mutant. *ECT1* transcript levels were increased in the *dag2-1* mutant compared to the wild-type (Fig. 4b). This result suggests that DAG2 can upregulate the expression of *ECT1*, which is consistent with the role of ECT1 in promoting seed germination.

We identified several *AAAAG* motifs in the *ECT1* promoter that had been previously recognized as binding motifs for Dof transcription factors in *Arabidopsis*. To confirm the direct association of DAG2 with the *ECT1* promoter, we performed EMSAs using biotin-labeled *ECT1* promoter fragments that include one *AAAAG* motif (Fig. 4c), with the same unlabeled fragments serving as competitors. DAG2 induced an upward shift of the *ECT1* promoter fragments (Supplementary Fig. 6a). Moreover, we mutated the probes used in the aforementioned experiment and found that DAG2 bound to the *ECT1* promoter fragment, whereas mutating the *AAAAG* motif abolished promoter association (Fig. 4d, e). These experimental results indicate that DAG2 binds to the *AAAAG* motif of *ECT1* promoter *in vitro*.

Collectively, the findings indicate that DAG2 is capable of directly interacting with the *ECT1* promoter, leading to an increasing its expression levels and ultimately facilitating seed germination.

### ECT1 enhance *DAG2* mRNA stabilization in *Arabidopsis*

DAG2 was found to be predicted as a direct target of ECT1 and capable of being modified by m^6^A from our comparison of the RIP-identified ECT1 target sites with the confident m^6^A methylation peaks from wild-type seedlings in the IPOP database (Miao et al., 2022; Huang et al., 2023). We assessed the binding ability of ECT1 to *DAG2* mRNA. To achieve this, we conducted RIP-qPCR using transgenic *Arabidopsis* plants expressing the ECT1-FLAG fusion protein. Remarkably, we observed a significant enrichment of *DAG2* transcripts within the immunoprecipitated fraction of the ECT1-FLAG plants compared to the wild-type control (Fig. 5a). These results support the notion that ECT1 can bind to *DAG2* transcripts.

**Fig. 5.**
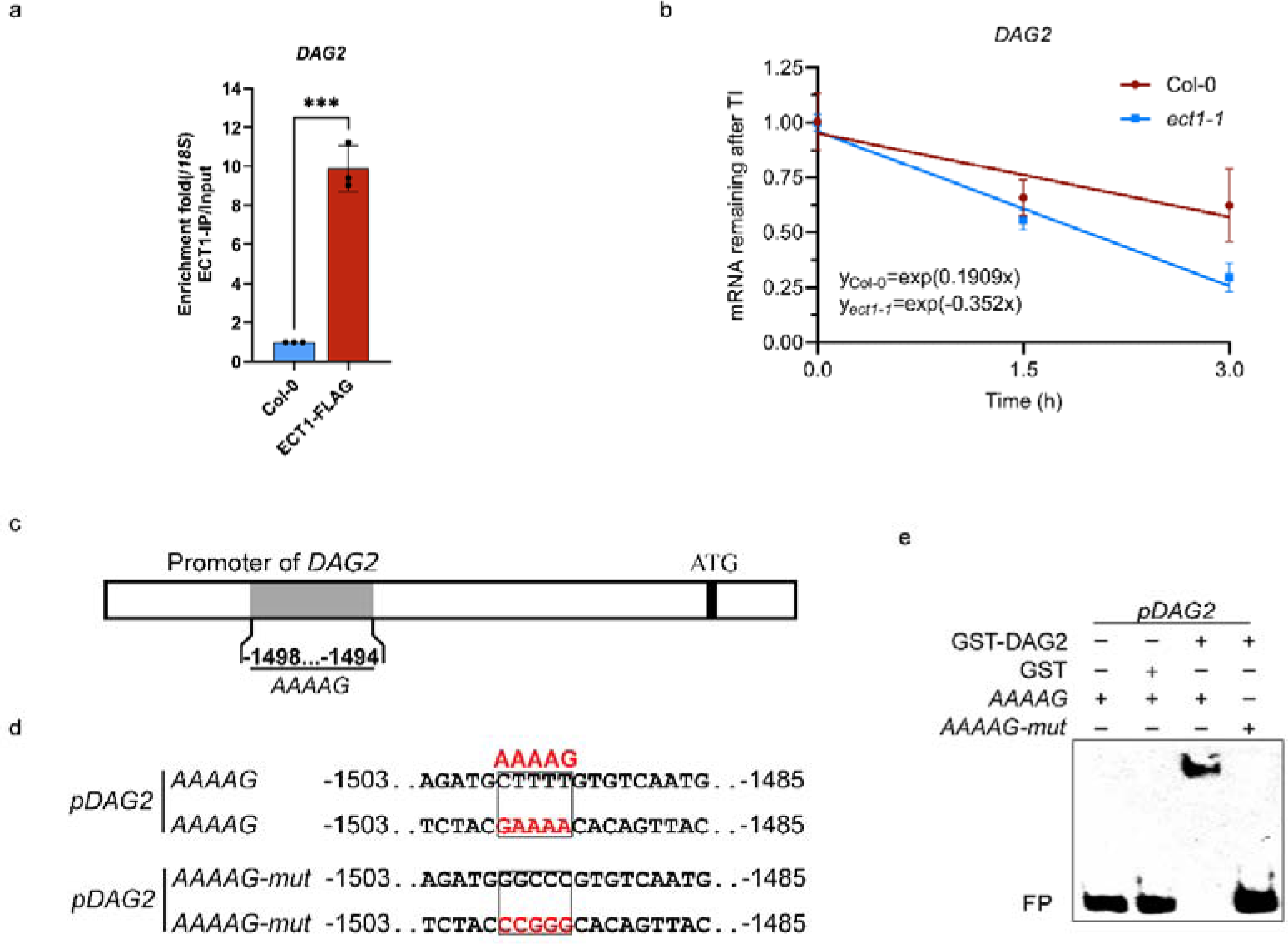
ECT1 interacts with *DAG2* transcripts and promotes *DAG2* stability. **(a)** RIP-qPCR assay. 4-day-old seedings of Col-0 *p35S: ECT1-FLAG* (line 8) and wild type Col-0 (negative control) were used in RIP assay. Chromatin was isolated and ECT1-FLAG bound RNA fragments were purified using anti-FLAG coupled magnetic beads. *DAG2* RNA fragments in the input and eluate fractions were detected using qPCR and specific primers. *18S* RNA was used for normalization. Values show enrichment of *DAG2* RNA fragments in the eluate fraction compared to the input fraction and are means of three replicates ± SD. Significant differences as determined by student’s t-test, ***: *p*<0.001. **(b)** mRNA stability assay. mRNA lifetimes of *DAG2* in Col-0 and *ect1-1* in 4-day-old seedlings. TI, transcription inhibition. *18S* RNA was used for normalization. Data are means ± SD for three replicates. **(c)** and **(d)** Schematic representation of the *DAG2* promoter. The promoter fragment used for EMSA is shown in (e). **(e)** Biotin-labelled *DAG2* −1485…−1503 promoter fragments with either a wild-type or mutated *AAAAG* motif were incubated with GST-DAG2, or GST alone (negative control). Samples were analyzed by native PAGE. The gel was blotted onto a nylon membrane and signals were detected by streptavidin-coupled horseradish peroxidase and ECL. FP, free probe.

Next, we performed transcription inhibition assays to elucidate further the functional significance of ECT1 in regulating *DAG2*. By comparing the decay rates of *DAG2* transcripts in wild-type and *ect1-1* mutant plants, we found a significantly higher decay rate for *DAG2* transcripts in the *ect1-1* mutant plants (Fig. 5b). This observation suggests that ECT1 appears to have a positive impact on the stability of *DAG2* transcripts, leading to a higher decay rate in *ect1-1* mutant.

Considering that ECT1 can regulate itself from multiple aspects, we also found *AAAAG* motif in the promoter of *DAG2* (Fig. 5c). Therefore, we investigated whether DAG2 can directly bind to its own promoter to activate its expression. Our EMSA results indicate that DAG2 can directly bind to the *DAG2* promoter (Fig. 5d, e; Supplementary Fig. 6b), suggesting that DGA2 probably, like ECT1, also directly regulates its own expression to enhance its functionality.

Taken together, our comprehensive investigation provides strong evidence for the direct binding of ECT1 to *DAG2* mRNA and is essential for governing the mRNA stability of *DAG2*. Additionally, we have also discovered that DAG2 can directly bind to its own promoter. These findings, combined with the results from the above experiments, indicate that on one hand, ECT1 functions as a positive regulator in the *DAG2* control pathway. On the other hand, both ECT1 and DAG2 not only regulate their own expression but also interactively control each other, thereby actively contributing to the regulation of seed germination mediated by gibberellins.

### ECT1 and DAG2 act independently to control the expression of *PHYB*

It has been previously reported that the inactivation of DAG2 impacts seed germination that is dependent on phyB (Santopolo et al., 2015). Our study intriguingly revealed that *PHYB* is a target of ECT1, as indicated by our RIP-seq data. Additionally, we identified the DAG2 binding site *AAAAG* in the promoter of *PHYB* (Fig. 6a). We then investigated whether PHYB is a common target for both ECT1 and DAG2.

**Fig. 6.**
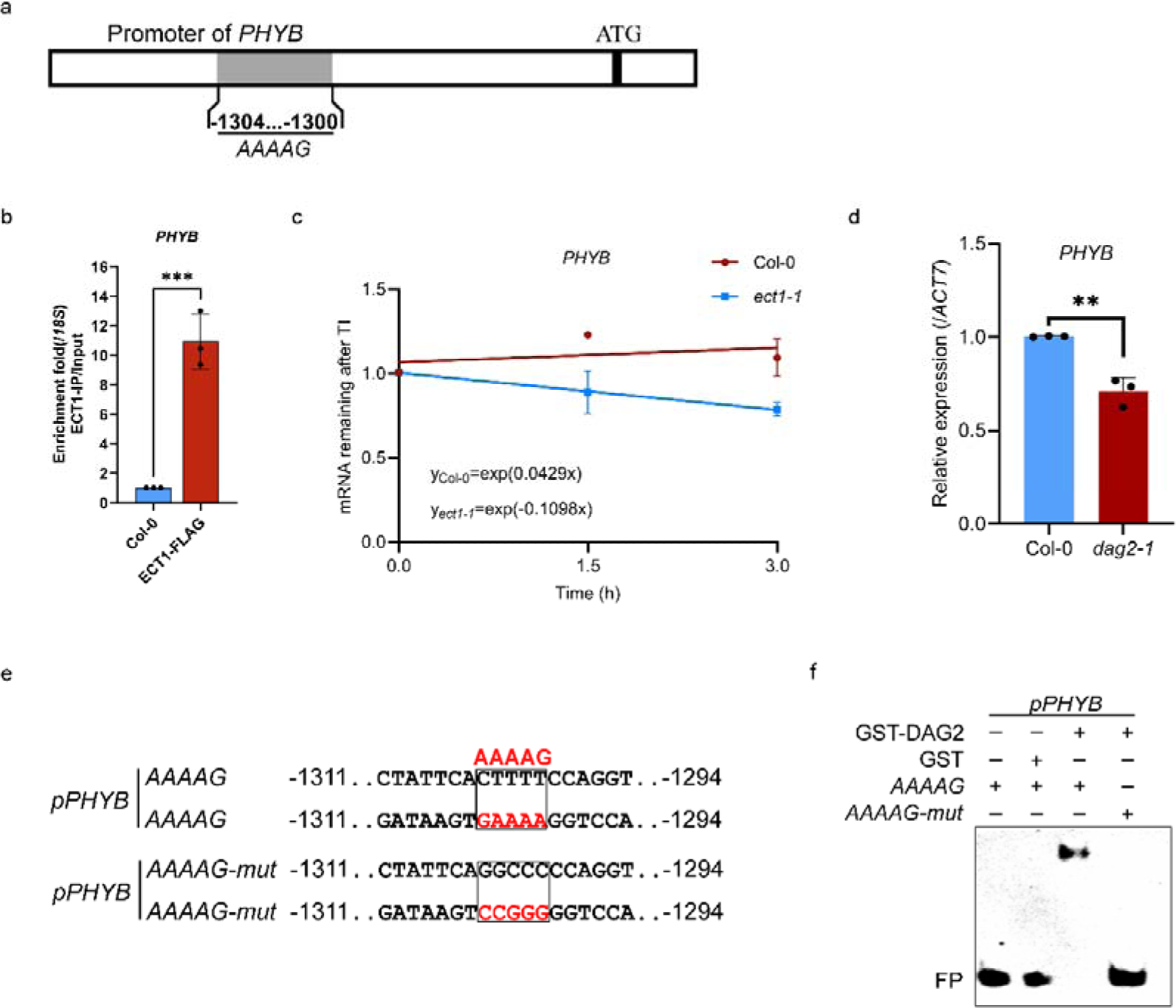
*PHYB* is a common target for both ECT1 and DAG2. **(a)** and **(e)** Schematic representation of the *PHYB* promoter. The promoter fragment used for EMSA is shown in (f). **(b)** RIP-qPCR assay. 4-day-old seedings of Col-0 *p35S: ECT1-FLAG* (line 8) and wild type Col-0 (negative control) were used in RIP assay. Chromatin was isolated and ECT1-FLAG bound RNA fragments were purified using anti-FLAG coupled magnetic beads. *PHYB* RNA fragments in the input and eluate fractions were detected using qPCR and specific primers. *18S* RNA was used for normalization. Values show enrichment of *DAG2* RNA fragments in the eluate fraction compared to the input fraction and are means of three replicates ± SD. Significant differences as determined by student’s t-test, **: P<0.01. **(c)** mRNA stability assay. mRNA lifetimes of *PHYB* in Col-0 and *ect1-1* in 4-day-old seedlings. TI, transcription inhibition. *Actin7* was used for normalization. Data are means ± SD for three replicates. **(d)** Comparison of *PHYB* mRNA expression in 4-day-old Col-0 and *dag2-1* seedlings. *Actin7* served as the reference gene for normalization. The presented data are the average of three replicates, with ± SD shown. Statistical significance was assessed using student’s t-test, with *** indicating *p*<0.001. **(f)** Biotin-labelled *PHYB* −1294…−1311 promoter fragments with either a wild-type or mutated *AAAAG* motif were incubated with GST-DAG2, or GST alone (negative control). Samples were analyzed by native PAGE. The gel was blotted onto a nylon membrane and signals were detected by streptavidin-coupled horseradish peroxidase and ECL. FP, free probe.

Initially, our RIP-qPCR experiments revealed that ECT1 has the capacity to complex with *PHYB*, which added to the evidence that *PHYB* is indeed a target of ECT1 (Fig. 6b). We next performed transcription inhibition assays to assess ECT1’s role in governing the stability of the *PHYB* transcript. We observed that the *ect1-1* mutant resulted in reduced stability of the *PHYB* transcript when compared to the wild-type (Fig. 6c). These findings implicate ECT1 as a positive factor in the recognition and stabilization of *PHYB*.

Given the presence of a DAG2 binding site in the *PHYB* promoter, we quantified *PHYB* expression in wild-type and *dag2-1* germinating seeds. The levels of *PHYB* transcripts were found to be reduced in *dag2-1* seeds compared to the wild type (Fig. 6d). To further substantiate the role of DAG2 in binding *PHYB* promoter, we conducted a EMSA experiments, which confirmed the direct binding of DAG2 to the *PHYB* promoter (Fig. 6 e, f; Supplementary Fig. 6c). These results suggest that DAG2 can directly bind to the *PHYB* promoter, which leads to an increase in *PHYB* expression and establishes a reinforcing feedback loop with phyB.

Overall, these findings suggests that both ECT1 and DAG2 are involved in the regulation of *PHYB* expression, with each acting as a positive regulator in their own distinct manner to facilitate the recognition and stabilization of *PHYB*.

## Discussion

The significance of m^6^A readers in plant growth and development is widely acknowledged. However, the precise identification of plant-specific m^6^A readers, responsible for interpreting and binding to m^6^A marks, has remained elusive (Wei et al., 2018; Song et al., 2023; Amara *et al*., 2023; Amara *et al*., 2024; Hou et al., 2021). Here, we have shown that ECT1 acts as an m^6^A reader protein, directly binding to the m^6^A site and playing a role in the regulation of seed germination in the presence of gibberellin. We have uncovered three distinct regulatory mechanisms of ECT1 that set it apart from other reported readers in plants. Firstly, ECT1 undergoes m^6^A modification itself to ensure its own mRNA stability, and its protein stability is further bolstered by the self-association of the ECT1 protein. Secondly, ECT1 engages in a regulatory feedback loop with DAG2, a known regulator of seed germination (Gualberti et al., 2002; Santopolo et al., 2015). ECT1 modulates the mRNA stability of *DAG2*, while DAG2 directly bind to the *ECT1* promoter to regulate its transcription. Lastly, both ECT1 and DAG2 operate independently to govern the expression of *PHYB*, which acts as an upstream regulator of DAG2 (Santopolo et al., 2015). ECT1 serves as a positive factor in the recognition and stabilization of *PHYB*, whereas DAG2 can bind directly to the *PHYB* promoter, resulting in an upregulating in *PHYB* expression.

We have provided evidence supporting ECT1 as an m^6^A reader protein through EMSA and mutational analysis (Fig. 1a, b), confirming its direct binding to m^6^A-modified RNAs. Additionally, RIP-seq analysis revealed the commonality of ECT1 with other m^6^A reader proteins (Fig. 1c-e). ECT1 belongs to the YTH family, which can be classified into YTHDC and YTHDF groups (Supplementary Fig. 7a; Zhang et al., 2010). In plants, YTH proteins are divided into five subfamilies: YTHDCA, YTHDCB, YTHDFA, YTHDFB, and YTHDFC (Supplementary Fig. 7a; Scutenaire et al., 2018). YTHDCA, YTHDCB, and YTHDFB are widely distributed in land plants, while YTHDFC is exclusive to seed plants, and YTHDFA is found only in flowering plants (Supplementary Fig. 7b). This classification offers valuable insights into the evolutionary diversification and functional roles of YTH proteins, with YTHDFA potentially representing a more recently emerged branch associated with functional diversification. ECT1 is one of the YTHDFA proteins (Supplementary Fig. 8). We observed that ECT1 directly binds to its own mRNA, which carries m^6^A modifications, resulting in the stabilization of its own transcripts (Fig. 2a, b). Furthermore, ECT1 undergoes self-interaction, forming an ECT1-ECT1 complex that enhances its protein stability (Fig. 2c-e). These findings provide valuable insights into the new functions and regulatory mechanisms of ECT1 as an m^6^A reader protein, shedding light on the functional and regulatory aspects of ECT1 in *Arabidopsis*, and contributing to our understanding of the m^6^A signaling pathway.

m^6^A plays a significant role in seed germination. The m^6^A “writer” components, including MTB and VIR, regulate seed germination in response to NaCl in an m^6^A-dependent manner (Hu et al., 2021). Several eraser proteins, namely ALKBH6, ALKBH8B, ALKBH9B, ALKBH9C, and ALKBH10B have been identified to play significant roles in ABA-regulated seed germination (Huong et al., 2020; Huong et al., 2022; Tang *et al*., 2022; Amara et al., 2022; Shoaib et al., 2021). Recent research also suggests that ECT2, ECT3, and ECT4 share similar functions in ABA-regulating seed germination (Song et al., 2023). Notably, our study uncovered a physiological role for ECT1 in gibberellin-induced germination. We observed that the *ect1-1* mutant exhibited hyposensitive to GA_3_ (Fig. 3a-c). This implies that ECT1 acts as a positive regulator in gibberellin-induced seed germination. In addition, we have demonstrated that GA_3_ positively regulates the expression of ECT1 at both the transcriptional and protein levels (Fig. 3d-e), confirming ECT1’s involvement in gibberellin-regulated seed germination. Seed germination is a crucial ecological and agronomic trait (Nonogaki et al., 2018; Li et al., 2022a). The presence of a single protein solely responsible for regulating this intricate process would pose a significant risk, as the loss of its function would halt the entire germination process. Functional redundancy among proteins serves as one of the ways to ensure the proper functioning of plants (Li et al., 2022b). Diverse “writer”, “eraser” and “reader” proteins can all perform functions during seed germination, and their modes of action are not entirely consistent, serving as one of the mechanisms to ensure the proper functioning of plants. In the future, studying how proteins from the same family balance the regulation of a particular trait will be a very intriguing research topic.

In addition, our study offers valuable perspectives on the regulatory mechanisms of ECT1 and its interaction with transcription factors DAG2. Utilizing a suite of ten bioinformatics resources, we were able to identify potential transcription factors that may govern *ECT1* expression (Supplementary Fig. 4a). Through an intersection with ECT1-RIP-seq data, we pinpointed four transcription factors, including DAG2, that potentially directly regulate *ECT1* expression and are under direct regulatory influence of ECT1. (Fig. 4a). Similar to ECT1, DAG2 also plays a positive role in the regulation of seed germination by gibberellins (Gualberti et al., 2002; Supplementary Fig. 5). The increased levels of *ECT1* in the *dag2-1* mutant (Fig. 4b), along with the direct binding of DAG2 to the *ECT1* promoter (Fig. 4c-d; Supplementary Fig. 6a), suggest it direct involvement in regulating *ECT1* expression. This demonstrate that DAG2 may regulates seed germination by directly controlling *ECT1*. Notably, the introduction of feedback regulation into the ECT1-controlled network adds a layer of complexity to the system. We have established that ECT1 plays a role in stabilizing *DAG2* mRNA, thereby negatively impacting the stability of *DAG2* transcripts (Fig. 5b, c). These findings enhance our understanding of the intricate regulatory networks involved in gene expression and emphasize the potential role of RNA modifications, such as m^6^A, in fine-tuning gene regulation. In addition, DAG2 is involved in the upregulation of gibberellin biosynthetic genes, specifically *AtGA3ox1* and *AtGA3ox2* (Santopolo et al., 2015). Our findings reveal that the expression level of ECT1 is enhanced by GA_3_ (Fig. 3d, e), which aligns with the conclusion that both ECT1 and DAG2 contribute positively to gibberellin-mediated seed germination. Future studies can explore the coordinated regulation of downstream gene expression and signaling pathways influenced by ECT1-DAG2 regulatory network, unraveling the broader implications of these mechanisms in plant growth and development.

Seed germination is regulated by several environmental factors, with light being a critical environmental cue that mediates seed germination in small-seeded plants such as *Arabidopsis* (Yang et al., 2020). Among photoreceptors, phyB is a primary phytochrome responsible for the regulation of seed germination and has been extensively studied (Kim et al., 2008). Previous research has indicated that DAG2 acts as a positive regulator of phyB-mediated seed germination (Santopolo et al., 2015). We have uncovered a refined layer of this regulatory mechanism, where DAG2 can feedback regulate phyB by directly binding to the *PHYB* promoter, thereby enhancing its expression (Fig. 6d-f; Supplementary Fig. 6c). This interplay is further enriched by the involvement of ECT1, which can form a regulatory loop with DAG2, also directly regulates *PHYB*, but through a different mechanism than DAG2. ECT1 regulates the stability of *PHYB* mRNA (Fig. 6b, c). The convergence of DAG2 and ECT1 on a common target, *PHYB*, yet via divergent pathways-one acting at the transcriptional level and the other at the post-transcriptional level-constitutes a robust and multifaceted strategy to ensure the precise expression of *PHYB*. The collaborative actions of DAG2 and ECT1 enhance the complexity of the molecular network that supports the phyB-gibberellin pathway, a critical pathway for triggering seed germination. Unlocking the intricacies of these interconnected mechanisms deepens our insight into the sophisticated processes governing plant responses to environmental cues, thereby broadening the scope for controlling seed germination.

In *Arabidopsis*, a total of thirteen YTH family proteins have been computationally identified as m^6^A reader proteins. Among these, ECT1 and ECT2 are classified under the DFA family, ECT8 under the DFC family, CPSF30-L under the DCA family, and ECT12 under the DCB family, while the other ECT proteins is yet to be strongly validated (Wei et al., 2018; Song et al., 2023; Amara *et al*., 2024; Hou et al., 2021). Our EMSA experiments revealed that ECT10 (belongs to DFB family) and ECT11 (belongs to DFC family) can specifically bind to m^6^A-modified RNA (Fig. 8), indicating their potential as additional m^6^A reader proteins. These findings lead us to speculate that all thirteen YTH proteins are involved in m^6^A recognition. ECT1, in particular, has been found to engage with its own mRNA, which has been marked by m^6^A modifications, pointing to a potential autoregulatory mechanism. The possibility that the other twelve YTH family members might also interact with *ECT1* in an m^6^A-dependent manner opens up a new avenue of investigation into the complex interplay between these m^6^A signaling regulators. Furthermore, the identification of DAG2 binding sites on the promoters of all thirteen YTH family proteins implies that DAG2 could play a central role in regulating the expression of these m^6^A readers, potentially influencing an array of biological processes, including seed germination and other critical traits. The discovery of these binding sites raises the intriguing possibility that DAG2 serves as a master regulator of the YTH family, orchestrating the epigenetic landscape that governs plant development and environmental responses. Unraveling the regulatory relationships between these different m^6^A reader proteins, their shared upstream and downstream genes, and the factors that modulate their activities will represent a fascinating and significant area of scientific inquiry.

In our comprehensive exploration of the m^6^A signaling cascade, we have uncovered ECT1 as a pivotal m^6^A reader protein, a discovery that marks a significant stride in our comprehension of this essential epigenetic pathway. Our research elucidates that ECT1 is a direct interactor with m^6^A-modified nucleotides, mediating a critical role in the regulation of seed germination in the presence of gibberellins. The stability of ECT1 is meticulously maintained through a dual mechanism: ECT1 undergoes m^6^A modification to ensure its own stability and the self-association of ECT1 protein enhances its stability. This multifaceted approach to ensuring ECT1’s stability is integral to its function as a regulator of gene expression. Our studies also reveal a sophisticated interplay between ECT1 and DAG2. ECT1 interacts with *DAG2* transcripts, thereby safeguarding the stability of *DAG2* mRNA. This interaction is reciprocated by DAG2, which binds to the promoter region of *ECT1*, thereby potentiating the expression of *ECT1*. This reciprocal regulation creates a feedback loop that fine-tunes the expression of both factors, thereby optimizing their combined cellular impact. The regulatory scope of ECT1 and DAG2 extends further to the positive regulation of *PHYB*, a central player in photomorphogenesis, through non-overlapping pathways. This independent yet convergent regulation of *PHYB* by ECT1 and DAG2 underscores the precision with which gene expression is controlled in response to environmental cues. These findings deepen our understanding of the complex regulatory networks governing m^6^A signaling and gene expression in plants, highlighting the fundamental importance of ECT1 in these processes.

## Plant Materials and Methods

### Plant materials and growth conditions

The *Arabidopsis* Columbia-0 (Col-0) ecotype was used as the wild-type in this study, the *ect1-1* mutant (SALK_073479C) and *dag2-1* mutant (SALK_201125C) in Col-0 was requested from AraShare (https://www.arashare.cn). Transgenic plants *p35S:ECT1-FLAG* was obtained by transforming plasmids into Col-0 plants. All *Arabidopsis* lines used in this work were grown in the greenhouse at 22°C under a 16 h day/8 h night cycle. Harvested seeds were dried at room temperature.

### Germination assay

Seeds from the same batch used in this experiment were harvested from plants that were cultivated under consistent conditions and simultaneously. As a result, the cumulative germination percentages for various treatments and genotypes within the same experimental framework are directly comparable, yet not across different experiments. For the germination assay, at least fifty seeds from each genotype were used, with the experiment conducted in three biological replicates. These seeds were left to acclimate at room temperature for four weeks before being used in the experiment.

To evaluate seed germination under the influence of gibberellic, the seeds were sterilized using chlorine gas for a period of forty minutes, after which they were transferred to a fume hood to remove any remaining chlorine gas. Following sterilization, the seeds were planted on a medium consisting of 1/2 MS/1.5% agar (1/2 MS Modified Medium, Coolaber, 1kg, China, Cat. No. PM10621-307) enriched with various concentrations of GA_3_ (Gibberellic Acid [GA3], Biotopped, 1g, China, CAS No. 77-06-5) as dictated in the Fig. 3a-c and Supplementary Fig. 6d, e. The plates were then incubated in the dark at 22℃ for four days before the germination process was assessed for completion. To confirm seed viability, each genotype was also planted on 1/2 MS/1.5% agar medium and subjected to white light exposure for four days.

### Protein expression and purification

To express GST-ECT1, GST-ECT1^W267A^, GST-ECT1^W324A^, GST-ECT1^W329A^, GST-DAG2, and GST proteins, the respective plasmids were transformed into *Escherichia. coli* strain BL21 (DE3) competent cells. These transformed cells were cultured in 15 mL LB medium containing 100 mg/mL ampicillin overnight at 37°C, and then transferred into 1 L LB medium (100 mg/mL ampicillin) at a dilution of 1:100. Upon reaching an OD_600_ of 0.6 to 0.8 at 37°C, the cells were induced with 0.2 mM IPTG (Isopropyl-1-thio-β-D-galactopyranoside). Following 20 hours of incubation at 16°C, the cell pellet from the 1 L culture was resuspended in 40 mL binding buffer (140 mM NaCl, 2.7 mM KCl, 10 mM Na_2_HPO4, 1.8 mM KH_2_PO_4_, pH 7.4) supplemented with 1 mM PMSF and 1 mM DTT. After sonication and centrifugation at 13,000 rpm for 30 minutes at 4°C, the supernatant was filtered through a 0.45 μM filter membrane. The filtered supernatant was then loaded onto a GST affinity column (Affinity chromatography column empty column tube, Beyotime, 6 mL, China, Cat no. FCL06; BeyoGold™GST-tag Purification Resin, Beyotime, 10 mL, China, Cat no. P2251) pre-equilibrated with equilibrium buffer (140 mM NaCl, 2.7 mM KCl, 10 mM Na_2_HPO_4_, 1.8 mM KH_2_PO_4_, pH 7.4) according to the manufacturer’s instructions. After washing the column with 40 mL of equilibrium buffer, the proteins were eluted with elution buffer (50 mM Tris-HCl pH 8.0, 10 mM reduced glutathione), mixed with glycerol to a final concentration of 20%, and stored at -80°C.

Similarly, MBP and MBP-ECT1 proteins were induced following the same procedure as the GST fusion proteins. The induced products were resuspended in 40 mL equilibrium buffer (20 mM Tris-HCl pH 7.4, 0.2 mM NaCl, 1 mM EDTA). After sonication and centrifugation, the supernatant was filtered through a 0.45 μM filter membrane and applied to amylose resin (NEW ENGLAND BioLabs, 10mL, USA, Cat. no. E8021S). The resin was washed with equilibrium buffer, and the proteins were eluted with equilibrium buffer supplemented with 10 mM maltose after washing the column with 30 mL equilibration buffer. The purified proteins were mixed with glycerol to a final concentration of 20% and stored at -80°C.

### Electrophoretic mobility shift assay

Electrophoretic mobility shift assays (EMSA) were conducted following the protocol outlined in Li et al. (2022b). The probes used in the EMSA experiments comprised directly synthesized RNA and DNA probes created by annealing complementary oligonucleotides. The m^6^A-modified RNAs and non-m^6^A-modified RNAs were each diluted to a concentration of 200 nM for the EMSA. For the DNA probes, annealing was achieved by heating to 95°C using primers listed in the supplementary Tab 4.

The EMSA reactions were carried out in a total volume of 20 μLof binding buffer (50 mM Tris-HCl pH 8.0, 750 mM KCl, 2.5 mM EDTA, 0.5% Triton-X 100, 62.5% glycerol, 1 mM DTT), with 0.4 nmol of probes labeled with 5’-biotin and 2 μg of purified proteins mixed and incubated at room temperature for 20 minutes. The protein-probe complexes were then separated by electrophoresis on a native 6% acrylamide gel for 1 hour at 120V in 0.5x Tris-Borate-EDTA buffer. Subsequently, the RNA/DNA was electroblotted onto a nitrocellulose membrane for 30 minutes at 380 mA and crosslinked to the membrane at 120 mJ*cm^-2^ using a UV crosslinker. Following this, the membrane was blocked in a 0.5% BSA buffer for 2 hours and then incubated with Stabilized Streptavidin-Horseradish Peroxidase Conjugate (Streptavidin/HRP, Solabio, 1 mL, China, Cat no. SE068), diluted 20,000 times in 0.5% BSA buffer, for 30 minutes at room temperature. The membrane was washed three times with PBST and then subjected to detection using a ClarityTM Western ECL Substrate kit (Clarity Western ECL Substrate, BIO-RAD, 500 mL, USA; Cat. #170-5061). All the experiments were conducted at least three replicates.

### RNA-immunoprecipitation assay

The RNA-immunoprecipitation assay (RIP) was conducted using 4-day-old seedlings of Col-0 and Col-0 *p35S:ECT1-FLAG* stable transgenic lines. A total of 2 grams of seedlings were submerged in 15 mL of PBS with 1.5% formaldehyde under vacuum conditions for 15 minutes to facilitate cross-linking. To stop the cross-linking reaction, 2.5 mL of 2 M glycine was added to the cross-linking reaction liquid and gently mixed, followed by vacuum treatment for 5 minutes. The seedlings were then transferred to a funnel fitted with paper to collect and washed with 200 mL of PBS. Subsequently, the seedlings were ground in liquid nitrogen and dissolved in 15 mL of nuclei extraction buffer (100 mM MOPS pH 7.6, 10 mM MgCl_2_, 5% Dextran T-40, 2.5% Ficoll 400, 0.5% [w/v] BSA [Fatty Acid & IgG Free, BioPremium, 20 g, Beyotime, Cat no. ST025], 10 mM DTT, 1× complete Protease Inhibitor Cocktail [Sigma-Aldrich, Germany, Cat no. 04693159001], 50 μM MG-132 [Sigma-Aldrich, 1 mg, Germany, Cat no. 474790], 0.4 M Sucrose, 1 mM PMSF, 40 U/mL RiboLock Rnase inhibitor [ThermoFisher, 2,500 units, USA, Cat no. EO0381]) for chromatin extraction. The mixture was then separated by four layers of miracloth to collect the liquid of nuclei preparation. After centrifugation at 13,000 rpm at 4°C for 15 minutes, the supernatant was transferred to a new tube and mixed with nuclei lysis buffer (50 mM Tris-HCl pH 8.0, 10 mM EDTA pH 8.0, 1% SDS, 1 mM PMSF) in a 1:1 ratio. The mixture was incubated on ice for 30 minutes. Then, 60 mL of anti-FLAG magnetic beads (Anti-FLAG^®^ M2 Magnetic Beads, MERCK, Germany, Cat no. M8823-1ML) were added to the mixture and incubated for 2 hours with gentle rocking at room temperature for chromatin immunoprecipitation. The immunoprecipitated complexes were eluted by adding 200 μL of nuclei lysis buffer heated to 65°C. To each sample, 1 μL of Proteinase K (BioLabs, 2 mL, USA, Cat no. P8107S) was added, and the samples were incubated at 37°C for 30 minutes for RNA reverse cross-linking. The RNA was extracted using the *SteadyPure* Plant RNA Extraction Kit (Accurate Biology, 50 rxns, China, Code No. AG21019) and used for RT-qPCR with gene-specific primers (Supplementary Tab. 3).

### RNA stability assay

The total RNAs used for the *in vitro* RNA stability assay were extracted from 4-day-old wild-type *Arabidopsis* seedlings. The assay reactions were carried out in a total volume of 300 μL of binding buffer (150 mM NaCl, 0.1% Nonidet P-40, 10 mM Tris-HCl, 0.5 mM DTT, pH 7.4) with actinomycin D added to a final concentration of 100 μM. For each reaction, 3 μg of GST-ECT1 and GST purified proteins were mixed and incubated at room temperature. At the beginning of the reaction, a 100 μL sample was taken from the reaction mixture and immediately frozen with nitrogen. The remaining samples were collected at 1.5 hours and 3 hours, and stored at -80°C. Prior to analysis, all samples were thawed on ice, and total RNAs were extracted using the *SteadyPure* Plant RNA Extraction Kit. Reverse transcription was performed using the 5x *Evo M-MLV* RT Master Mix (*Evo M-MLV* RT Premix for qPCR, Accurate Biology, China, Cat no. AG11706) immediately after extraction. The expression of *ECT1* in different samples was detected using qPCR with primers listed in the supplementary Tab. 3.

For the *in vivo* RNA stability assay, a modified version of a previously described assay (Song et al., 2023) was used. 4-day-old wild-type and *ect1-1 Arabidopsis* seedlings grown on filter paper were transferred to a 15 mL centrifuge tube containing 4 mL of ddH_2_O with a final concentration of 100 μM actinomycin D. After 1 hour of infiltration, the seedlings were harvested and used as the 0-hour time control. Subsequent samples were harvested every 1.5 hours and immediately frozen in liquid nitrogen. RNA isolation and RT-qPCR analysis were performed to quantify the remaining mRNA levels, with *18S* RNA serving as the internal control. The experiment was conducted with 3 biological replicates and three technical replicates per sample.

### Protein extraction and immunoblotting

The 4-day-old wild-type and ECT1OE-8 plants were used for protein extraction in this study. To extract proteins, 100 mg of seedlings were ground to a fine powder using a high-throughput tissue grinding machine and 200 μL of extraction buffer (65 mM Tris, 4 M urea, 10% [v/v] glycine, 10 µM DTT, 1x protease inhibitor cocktail [Roche], pH 7.3) was added. After vertexing and shaking, the samples were centrifuged at 12,000 rpm for 10 min at 4°C to remove the supernatant, and 4× SDS protein loading buffer (containing 1 M Tris-HCl pH 8.0, 1 M DTT, 10% SDS, 40% glycine, 0.4% bromophenol blue) was added to the supernatant and heated at 98°C for 10 min.

For the protein expression assay experiments. The extract proteins samples were separated by electrophoresis in 12% SDS-PAGE and then transferred to a PVDF membrane. The membrane was blocked in 25 mL blocking buffer (5% milk in PBS) for 1 hour at room temperature, then washed three times with 1xPBS for 5 minutes each, and incubated with the appropriate antibody. The membrane was subjected to a 2-hour incubation at room temperature with either a FLAG tag-specific mouse monoclonal antibody (diluted 1:1000, FLAG Tag Mouse Monoclonal Antibody [HRP Conjugated], Beyotime, 50 μL, China, Cat no. AF2855) or a monoclonal antibody against mouse β-actin (diluted 1:5000, LABLEAD, 100 μL, China, Cat no. BP0101). The membrane was washed three times with 1x PBST buffer for 5 min each time, and then incubated for 1 h at room temperature with the horseradish peroxidase (HRP)-linked goat-conjugated mouse IgG secondary antibody (1:2500, Beyotime, 1 mL, China, Cat no. A0216). After washing the membrane twice with PBST and once with PBS, the Clarity^TM^ Western ECL Substrate was applied to the membrane and the luminescence signal was detected with a CCD camera.

To assess protein stability, proteins extracted from 4-day-old ECT1OE-8 and wild-type seedlings were placed in centrifuge tubes. Equimolar amounts of GST-ECT1 and GST control proteins were added to the samples, which were then gently mixed and left to incubate at room temperature for 2 hours. Following incubation, 4x SDS sample loading buffer was introduced to the mixture, which was subsequently heated to 98°C for 10 minutes to denature the proteins. The impact of GST-ECT1 on the stability of the ECT1 protein was then analyzed using western blotting, with the same protocol and antibodies employed as in the preceding assay. The experiment was conducted with three replicates.

### Yeast two hybrid (Y2H) assays

The Y2H assay was performed according to the protocol outlined by Li et al. (2022b). In summary, the CDS of *ECT1* was cloned into the *pGBKT7* vector (Sheerin et al., 2015), while the CDS of *ECT1/2/3/4* were cloned into the *pGADT7* (Sheerin et al., 2015) vector individually. The bait and prey vectors were co-transformed into the yeast strain AH109 using the Yeast Two-Hybrid Interaction Proving Kit (Coolaber, China, Cat no. YH2021). The transformed yeast cells were growing on SD medium lacking leucine and tryptophan (SD -Leu/-Trp) at 30°C. The transformants were then cultivated at 30°C with shaking at 200 rpm until the OD_600_ reached 0.6. Subsequently, they were grown on SD medium lacking leucine, tryptophan, and histidine (SD-Leu/-Trp/-His) supplemented with 3-amino-1,2,4-triazole (3-AT). The diluted transformants (10, 100, and 1000-fold dilutions with ddH_2_O) were plated on the selective medium to assess the interaction.

### Pull down assay

The pull down reactions were carried out in a total volume of 1 mL of pull down buffer (50 mM Tris-HCl, 200 mM NaCl, 1 mM EDTA, 1% NP-40, 1 mM DTT, 10 mM MgCl_2_, pH 8.0). 2 μg of purified proteins with GST- and MBP-tags were mixed and incubated under gentle shaking for 2 hours on ice. A 100 μL aliquot of the mixture was saved as the input and boiled with 25 μL of 4x SDS PAGE buffer (1 M Tris-HCl pH 8.0, 1 M DTT, 10% SDS, 40% glycine, 0.4% bromophenol blue). The remaining protein mixture was mixed with GST-tag purification resin (BeyoGold™GST-tag Purification Resin, Beyotime, 10 mL, China, Cat no. P2251) and incubated under gentle shaking for 2 hours on ice. The resin was washed three times with PBS buffer (140 mM NaCl, 2.7 mM KCl, 10 mM Na_2_HPO4, 1.8 mM KH_2_PO_4_, pH 7.3), and the proteins were eluted by boiling with 4x SDS PAGE buffer. The samples were separated by SDS-PAGE at 120 V and transferred onto a PVDF membrane at 200 mA for 2 hours. The membrane was blocked with 5% blocking buffer (5% nonfat-dried milk dissolved in PBST) under gentle shaking for 2 hours at room temperature. After washing twice with PBST, the membrane was incubated separately with anti-GST antibody (GST Tag Mouse Monoclonal Antibody [HRP Conjugated], Beyotime, 50 μL, China, Cat no. AF2891) and anti-MBP antibody (MBP Tag Mouse Monoclonal Antibody [HRP Conjugated], Beyotime, 50 μL, China, Cat no. AF2915) at a dilution of 1:2000 for 2 hours at room temperature. The membrane was washed three times with PBST for 10 minutes each and then the luminescence signals were detected using the Clarity Western ECL Substrate. The experiment was conducted with three replicates.

### Quantification of transcript levels by RT-qPCR

For the gene expression analysis, total RNA was extracted from 4-day-old seedlings using a commercial RNA extraction kit (Accurate Biology, 50 rxns, China, Code No. AG21019), and cDNA was synthesized according to the protocol of the 5x *Evo M-MLV* RT Master Mix (*Evo M-MLV* RT Premix for qPCR, Accurate Biology, China, Cat no. AG11706). The mRNA levels of the target genes were measured using the SYBR Green Premix Pro Taq HS qPCR Kit (Accurate Biology, China, Cat no. AG11701) and gene-specific primers (refer to Supplementary Tab.3). The qPCR was performed on a LightCycler 96 instrument (Roche) with the following cycling conditions: initial denaturation at 95°C for 30 seconds, followed by 40 cycles of denaturation at 95°C for 5 seconds, and annealing/extension at 60°C for 30 seconds. The relative expression levels were calculated using the 2^-ΔΔCT^ method. Statistical significance was determined using a t-test. The experiments were performed with three biological replicates and three technical replicates for each sample.

### Phylogenetic analysis

The protein sequences in FASTA format and genome annotations in GFF3 format for 26 species were downloaded from public databases, including Ensembl (https://www.ensembl.org/index.html), Ensembl Plants (https://plants.ensembl.org), Phytozome (https://phytozome.jgi.doe.gov/pz/portal.html), and the National Center for Biotechnology Information (NCBI, https://www.ncbi.nlm.nih.gov). These species included two animals, two red algae, two green algae, two mosses, two ferns, two basal angiosperms, seven monocots, and seven dicots (Supplementary Tab. 8). To identify the YTH family genes, a hidden Markov model (HMM) profile (PF04146) from the InterPro database (https://www.ebi.ac.uk/interpro) was used with an E-value threshold of 1e-5. All candidate genes were further examined using the Conserved Domain Database (CDD, https://www.ncbi.nlm.nih.gov/Structure/cdd/wrpsb.cgi). The resulting YTH candidate protein sequences were input in MAFFT version 7.520 (Katoh et al., 2002) for multiple sequence alignment. Using the YTH protein of *Chlamydomonas reinhardtii* as an outgroup, a phylogenetic tree analysis of YTH proteins was conducted using the maximum likelihood method with IQ-TREE version 2.2.0 (Minh et al., 2020). The bootstrap value was set to 1000.The species phylogenetic tree was downloaded from TimeTree (https://timetree.org). The phylogenetic trees were visualized using the R package ggtree version 3.8.2 (Yu et al., 2017) and Interactive Tree Of Life version 6 (iTOL v6) (Letunic and Bork, 2021). Following with these methods, the YTH protein sequences were aligned and phylogenetic trees were constructed for understanding the diversity of YTH protein in Arabidopsis thaliana. The multiple sequence alignment results were visualized using ESPript version 3.0.10 (Robert and Gouet, 2014), with a focus on highlighting conserved tryptophan residues.

### Regulation of network construction

The transcription factor regulatory network for ECT1 using three different approaches: collecting published regulatory networks, constructing sequence-based regulatory networks, and developing expression-based regulatory networks. Initially, we extracted upstream transcription factors of ECT1 from iRegNet (Shim et al., 2021) and AtRegNet (Palaniswamy et al., 2006), as well as from gene regulatory networks reported in previous studies (Chen et al., 2018; De Clercq et al., 2021). Additionally, we employed Random Forest-based regression models to predict potential interactions between transcription factors and ECT1 using the GENIE3 version 1.18.0 (Gene Network Inference with Ensemble of trees) algorithm (Huynh-Thu et al., 2010), utilizing TPM expression values of wild-type *Arabidopsis thaliana* Col-0 from 3131 samples in the *Arabidopsis* RNA-Seq Database (ARS) (Zhang et al., 2020). Furthermore, we predicted potential transcription factor binding sites within the promoter region of *ECT1* using PlantTFDB v5.0 (Jin et al., 2017), PlantPAN 4.0 (Chow et al., 2019), CIS-BP (Weirauch et al., 2014), JASPAR 2024 (Khan et al., 2018) and deepTFBS (https://github.com/cma2015/deepTFBS). Finally, transcription factors that were supported with four or more of the mentioned sources were retained for constructing ECT1-related gene regulatory network. The regulatory network was visualized using Cytoscape version 3.10.1 (Shannon et al., 2003).

### Homology modeling

The amino acid sequence of *ECT1* was submitted to the Swiss-Model server (https://swissmodel.expasy.org) to generate its three-dimensional structures. The chain of a representative YTHDF2-m^6^A complex (PDB ID: 4RDN) (Li et al., 2014) was used as the structural template. Molecular docking analyses were performed using the GOLD software suite v5.3 (Verdonk et al., 2003), and the resulting structures were visualized using PyMol version 2.5.4 (https://pymol.org).

### RIP-seq and Data Analysis

The RIP-seq work was carried out by LC-Bio Technologies (LC-Bio Technologies (Hangzhou) CO. LTD, Hangzhou, China). Seedlings of ECT1-FLAG (ECT1OE-8), at 4 days old and weighing 1 g, which had been cultivated on 1/2 MS agar plates, were collected and utilized for the RIP-seq experiment.

Briefly, harvest an appropriate quantity of tissue culture cells. To prepare the cell lysate, resuspend the final cell pellet in an equal volume of polysome lysis buffer, which consists of 100 mM KCl, 5 mM MgCl_2_, 10 mM HEPES (pH 7.2), 0.5% NP-40, 1 mM DTT, RNase inhibitors, and protease inhibitors. Incubate the mixture on ice for 5 minutes, followed by centrifugation at 4°C and 15,000 x g for 15 minutes to pellet cellular debris. After centrifugation, carefully collect 40 μL of the supernatant for RNA extraction to serve as the input sample. The remaining supernatant is reserved for the immunoprecipitation step. For the Antibody Coating of Protein A/G Beads. Begin by washing 100 μL of protein A/G magnetic beads with NT2 buffer, comprising 50 mM Tris-HCl (pH 7.4), 150 mM NaCl, 1 mM MgCl_2_, and 0.05% NP-40, a total of three times. Subsequently, resuspend the washed beads in 850 μL of the immunoprecipitation reaction solution, which includes 200 units of an RNase inhibitor, 400 μM RVC, 10 μL of 100 mM DTT, 30 μL of 0.5 mM EDTA, and 800 μL of cold NT2 buffer. Incubate the cleared cell lysates with this reaction solution at 4°C for 4 hours on a tube rotator. Proceed to wash the beads five times with 1 mL of ice-cold NT2 buffer. To isolate total RNA, add TRIzol to both the input and immunoprecipitation (IP) samples. Evaluate RNA quality control on a Qubit. Prepare the sample libraries using the SMART-Seq v4 Ultra Low Input RNA Kit (Takara Clontech Kit Cat# 63488) and sequence them on a NovaSeq 6000 (Illumina).

The quality of RIP-seq data (raw reads) was examined using the fastp software (version 0.20.1). To generate clean reads, adapter sequences and low-quality reads w ere removed with "-5 -3 -r --detect_adapter_for_pe -n_base_limit 0 -average_qual 20 " parameters. The clean reads were further aligned to the reference genome of Arabidopsis (TAIR10) with hisat2 (version 2.1.0). Peak calling was then performedusing ex omepeak2 (version 4.4). Gene annotation of the *Arabidopsis*genome was obtained from "https://www.arabidopsis.org/download_files/Genes/Araport11_genome_release/Araport11_GTF_genes_transposons.Mar202021.gtf.gz". Stringtie was used to quantify gene expression of input and filter out false positive peaks with lowly expressed target genes. HOMER (http://homer.ucsd.edu/homer/download.html, v4.11.1) was used to detect enriched motifs from ECT1-binding peaks. The top rank motif was visualized using ggseqlogo R package.

## Author contributions

Conceptualization: ZL, CM. Investigation: ZL, YM, WS, YB, PD, YQ, CJ, BL, CM. Visualization: ZL. Supervision: ZL, CM. Funding acquisition: ZL, CM. Writing—original draft: ZL, CM. Writing—review and editing: ZL, YM, WS, YB, PD, YQ, CJ, BL, CM.

---

**Supplementary Fig. 1.**
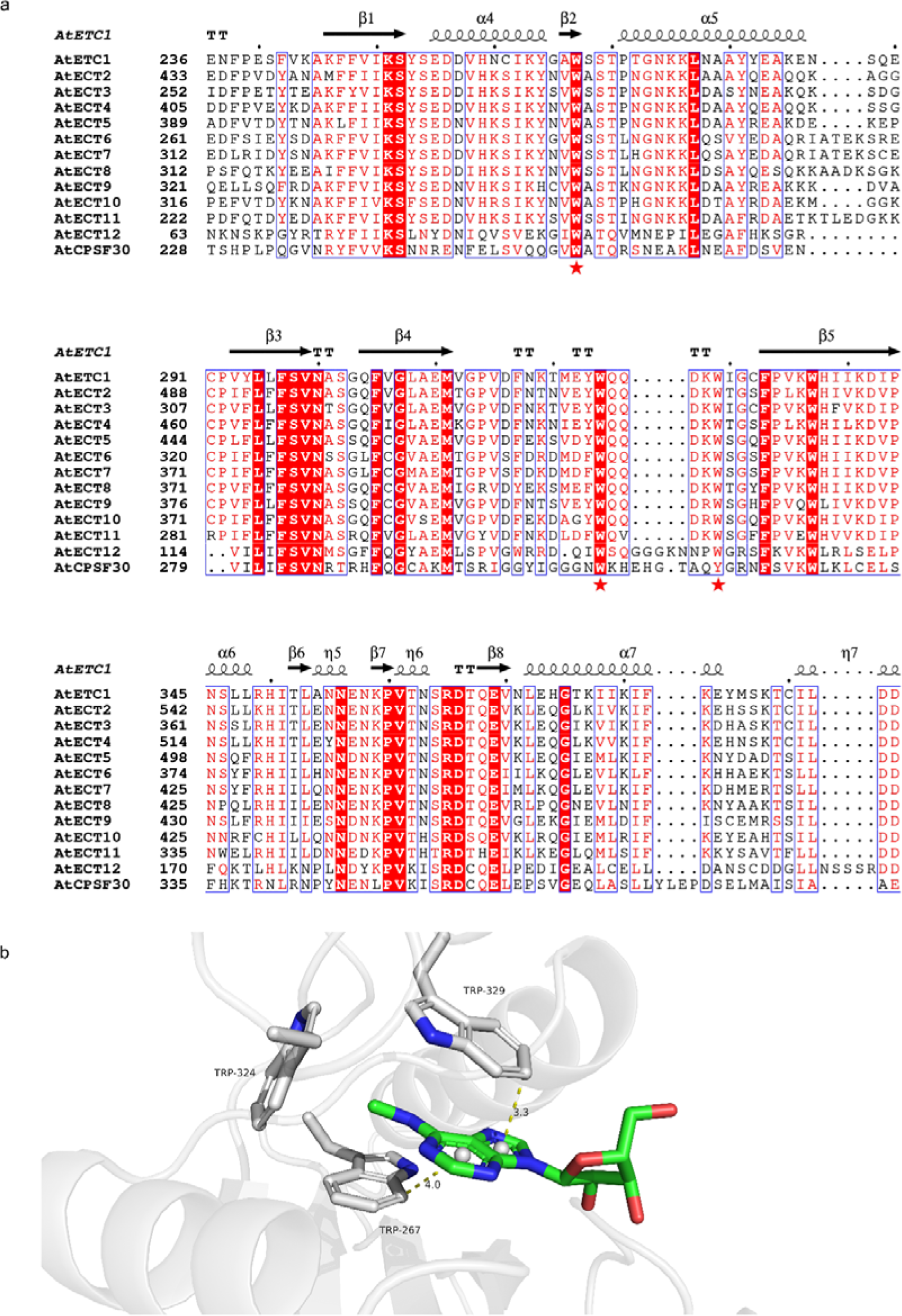
Sequence and structural analysis of the ECT1 Protein. **(a)** Multiple sequence alignment of the YTH domain of ECT proteins. The secondary structure elements of ECTs are indicated, and amino acids are colored according to the level of sequence conservation: red letters, similar residues; red boxes, identical residues. Amino acids that form the methyl-interacting aromatic cage are marked with red stars. **(b)** Model of the YTH domain of the ECT1 protein generated by the homology modeling server SWISS-MODEL. Trp267, Trp324, and Trp329 represent the three conserved Trp residues in ECT1, with another highlighted region representing the m^6^A-modified adenosine.

**Supplementary Fig. 2.**
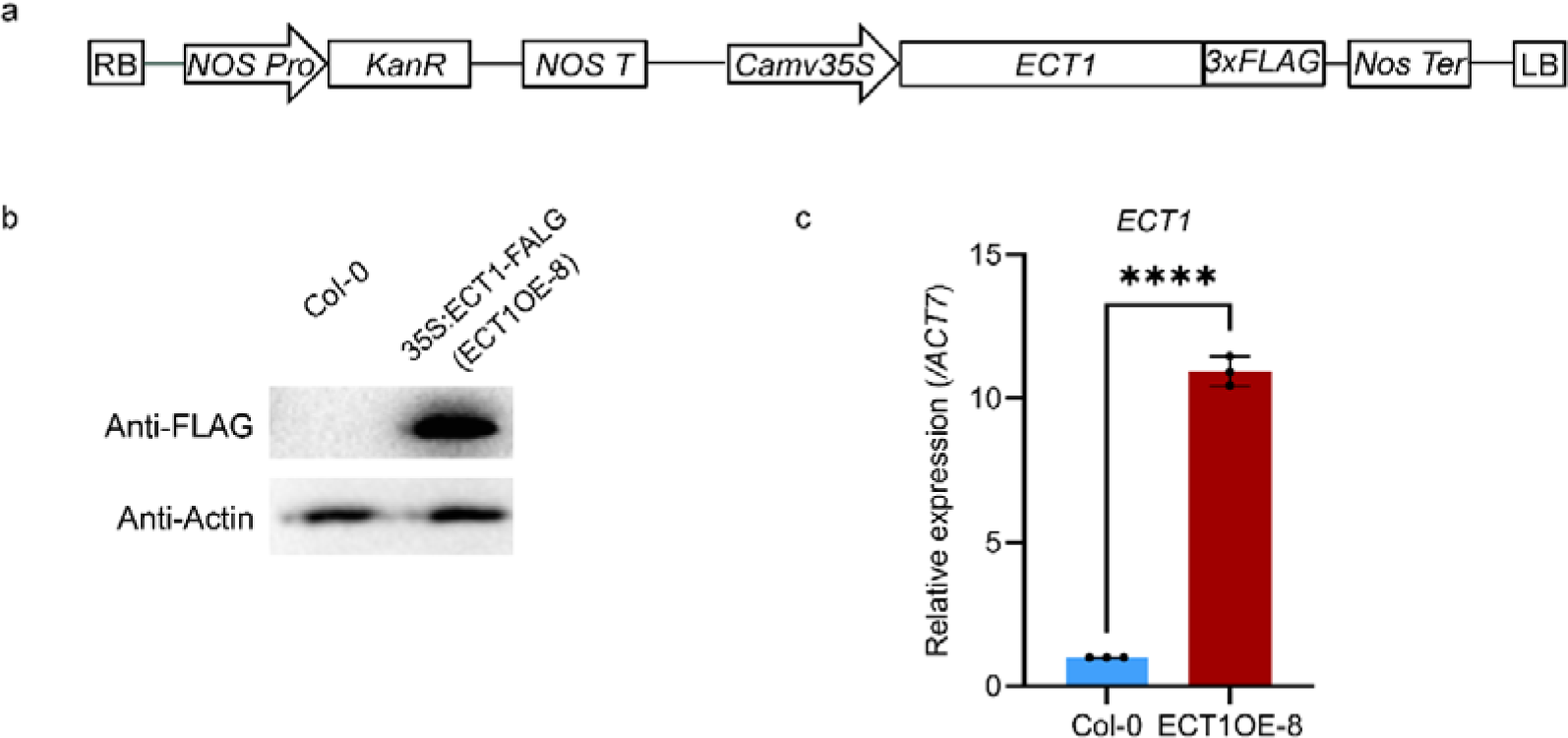
Identification of ECT1 overexpression line. **(a)** Schematic diagram of the construction of the *35S*:*ECT1*-*3×Flag* transgenic overexpression vector. **(b)** ECT1-FLAG protein expression in the ECT1 overexpressing line. Proteins were extracted from 4-day-old Col-0 and Col-0 *p35S: ECT1-FLAG* (line 8) seedlings and subjected to SDS-PAGE and immunoblot analysis using an anti-Flag antibody; anti-Actin was utilized as a loading control to detect ACTIN. Col-0 seeds served as a negative control. **(c)** Comparison of *ECT1* mRNA expression in 4-day-old Col-0 and Col-0 *p35S: ECT1-FLAG* (line 8) seedlings. *Actin7* served as the reference gene for normalization. The presented data are the average of three replicates, with ± SD shown. Significant differences as determined by student’s t-test, ***: *p*<0.0001.

**Supplementary Fig. 3.**
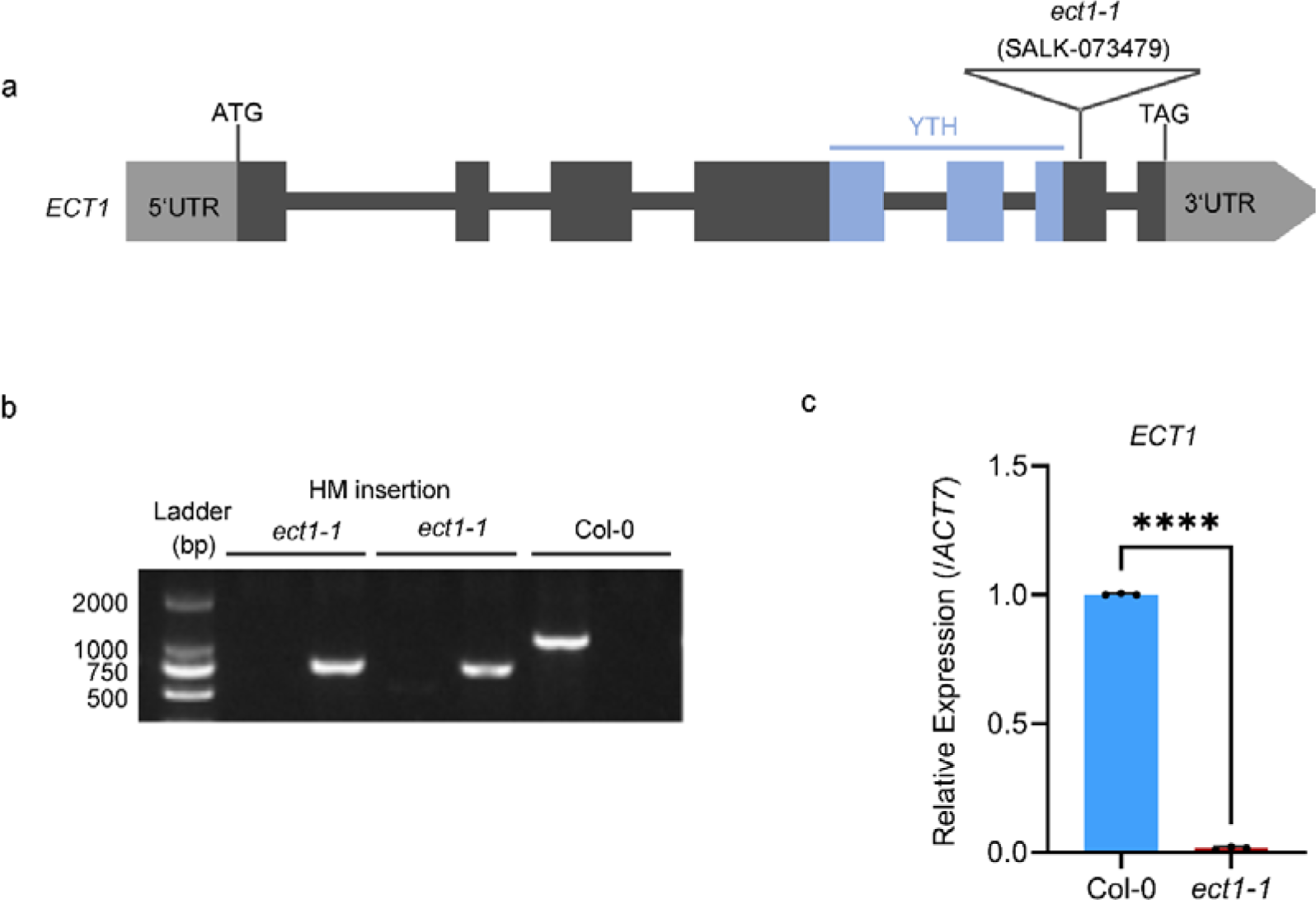
Identification of the *ECT1* T-DNA insertion line. **(a)** Diagrams of the T-DNA insertion sites within the *ECT1* locus in the *ect1-1* line. The gray box denotes the UTR, the black box signifies the exon, the black line denotes the intron, and the blue box encodes the YTH domain of ECT1. **(b)** PCR assay for the detection of *ECT1* CDS and a partial T-DNA fragment in genomic DNA (gDNA) from Col-0 and *ect1-1* mutant plants. **(c)** RT-qPCR analysis for the quantification of *ECT1* transcript levels in 4-day-old wild-type and *ect1-1* mutant seedlings. *Actin7* was used as an internal control. Values are means of three replicates ± SD. Significant differences as determined by student’s t-test, ***: *p*<0.0001.

**Supplementary Fig. 4.**
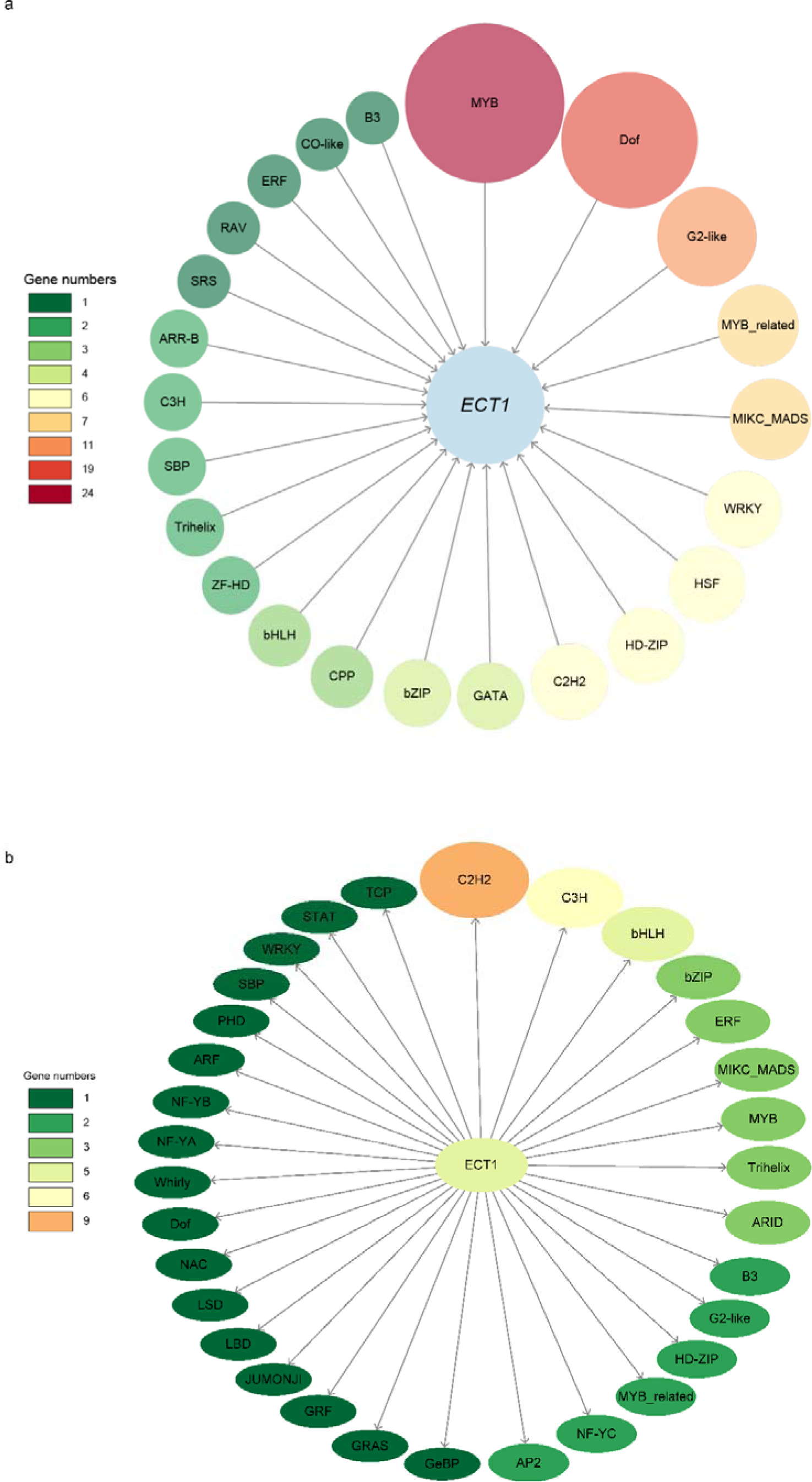
Identification of upstream and downstream regulatory genes linked to ECT1. (a) Diagram illustrating the putative upstream transcription factors of *ECT1* gene. (b) Diagram illustrating the downstream transcription factors targeted by the protein ECT1.

**Supplementary Fig. 5.**
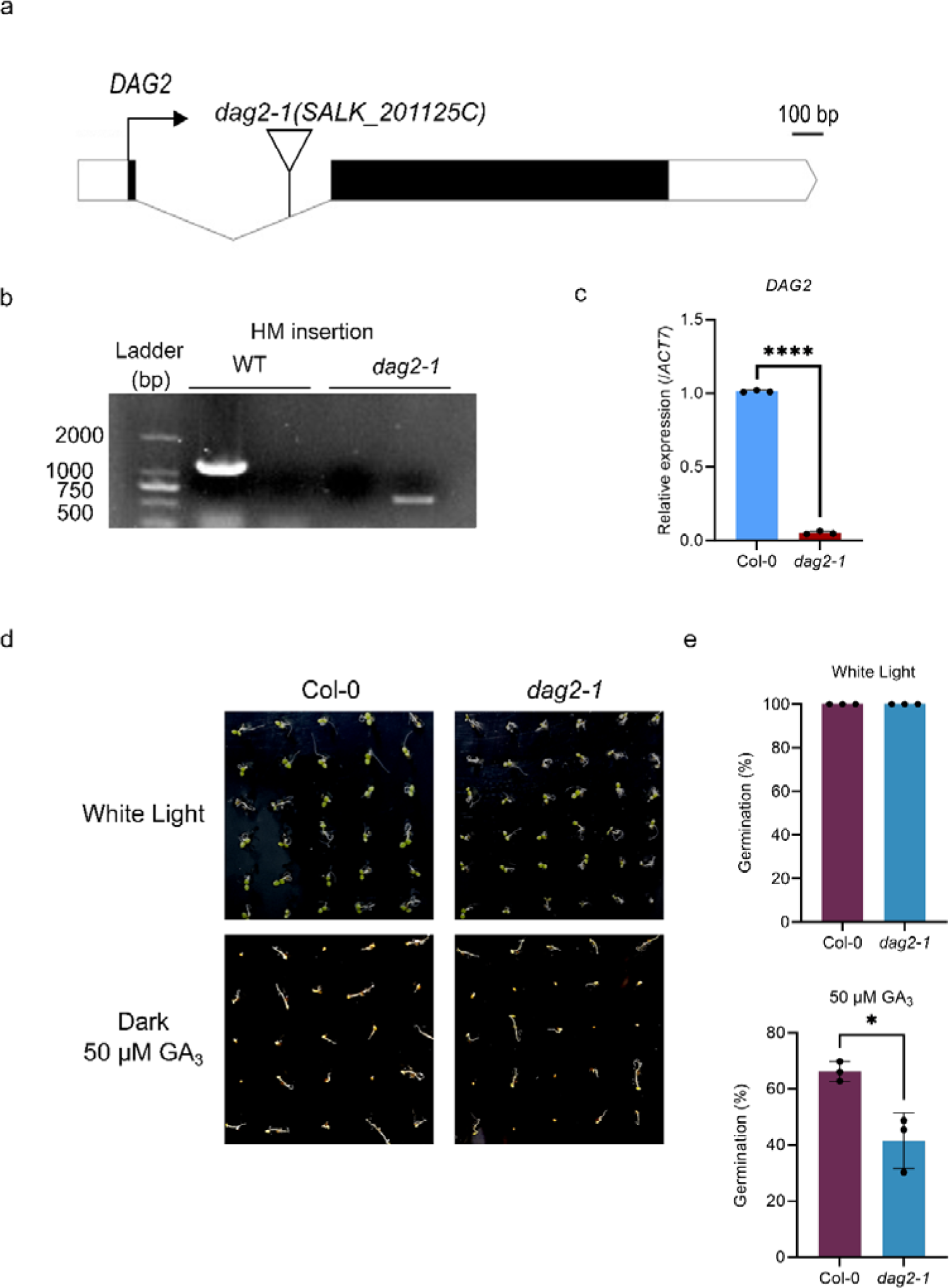
Identification of the *DAG2* T-DNA insertion line. **(a)** Diagrams of the T-DNA insertion sites within the *DAG2* locus in the *dag2-1* line. The white box denotes the UTR, the black box signifies the exon, the line denotes the intron. **(b)** PCR assay for the detection of *DAG2* CDS and a partial T-DNA fragment in gDNA from Col-0 and *dag2-1* mutant plants. **(c)** RT-qPCR analysis for the quantification of *DAG2* transcript levels in 4-day-old wild-type and *dag2-1* mutant seedlings. *Actin7* was used as an internal control. Values are means of three replicates ± SD. Significant differences as determined by student’s t-test, ***: *p*<0.0001. **(d)** Phenotypic assessment of the GA response in Col-0 and *dag2-1* seeds cultivated on 1/2 MS medium supplemented with 0 or 50 μM GA_3_, under either white light or dark conditions. Photographs representing the growth were captured four days after sowing. **(e)** Analysis of germination percentages in Col-0 and *dag2-1* mutants treated with 0 or 50 μM GA_3_, in either white light or dark conditions. Data from three replicates were averaged. The bar graph represents the average cumulative germination rates from three trials, with error bars indicating ± SD. Significant differences as determined by student’s t-test, *: *p*<0.05.

**Supplementary Fig. 6.**
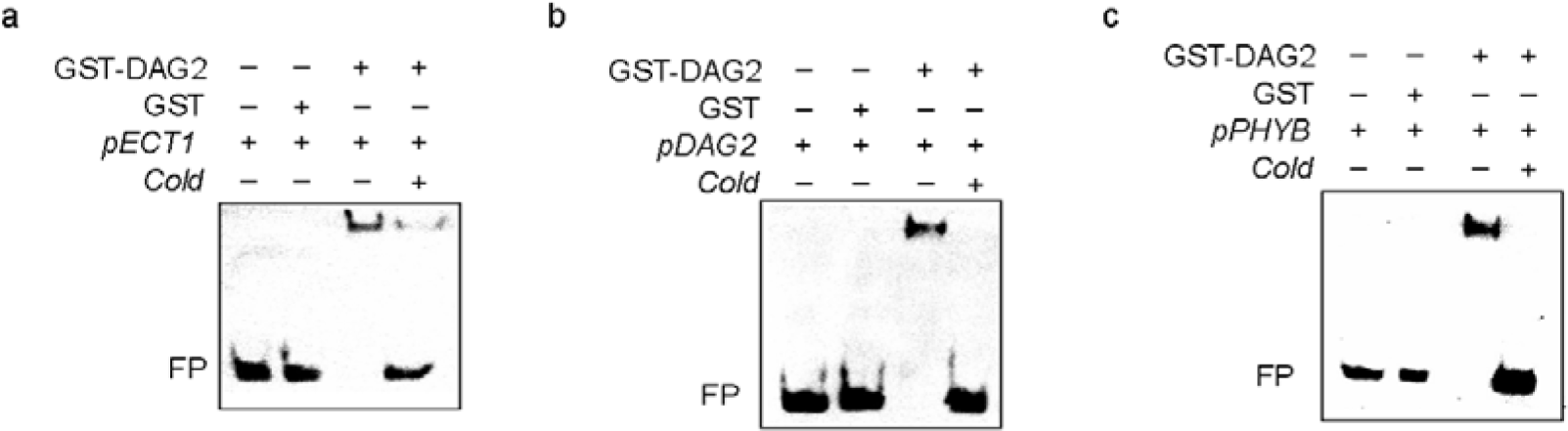
DAG2 interact with the *ECT1*, *DAG2* and *PHYB* promoters. **(a), (b), and (c)** Biotin-labelled *ECT1*, *DAG2* and *PHYB* promoter fragments (*pECT1* −810…−827, *pDAG2* −1485…−1503, and *pPHYB* −1294…−1311) were incubated with GST-DAG2, or GST alone (negative control). The gel was blotted onto a nylon membrane and signals were detected by streptavidin-coupled horseradish peroxidase and ECL. FP, free probe. Cold, excess of the DNA fragment, but unlabeled. In the experimental groups containing "cold" during the process, "cold" is added first to allow it to fully bind to GST-DAG2, and then the tested promoter is added ten minutes later.

**Supplementary Fig. 7.**
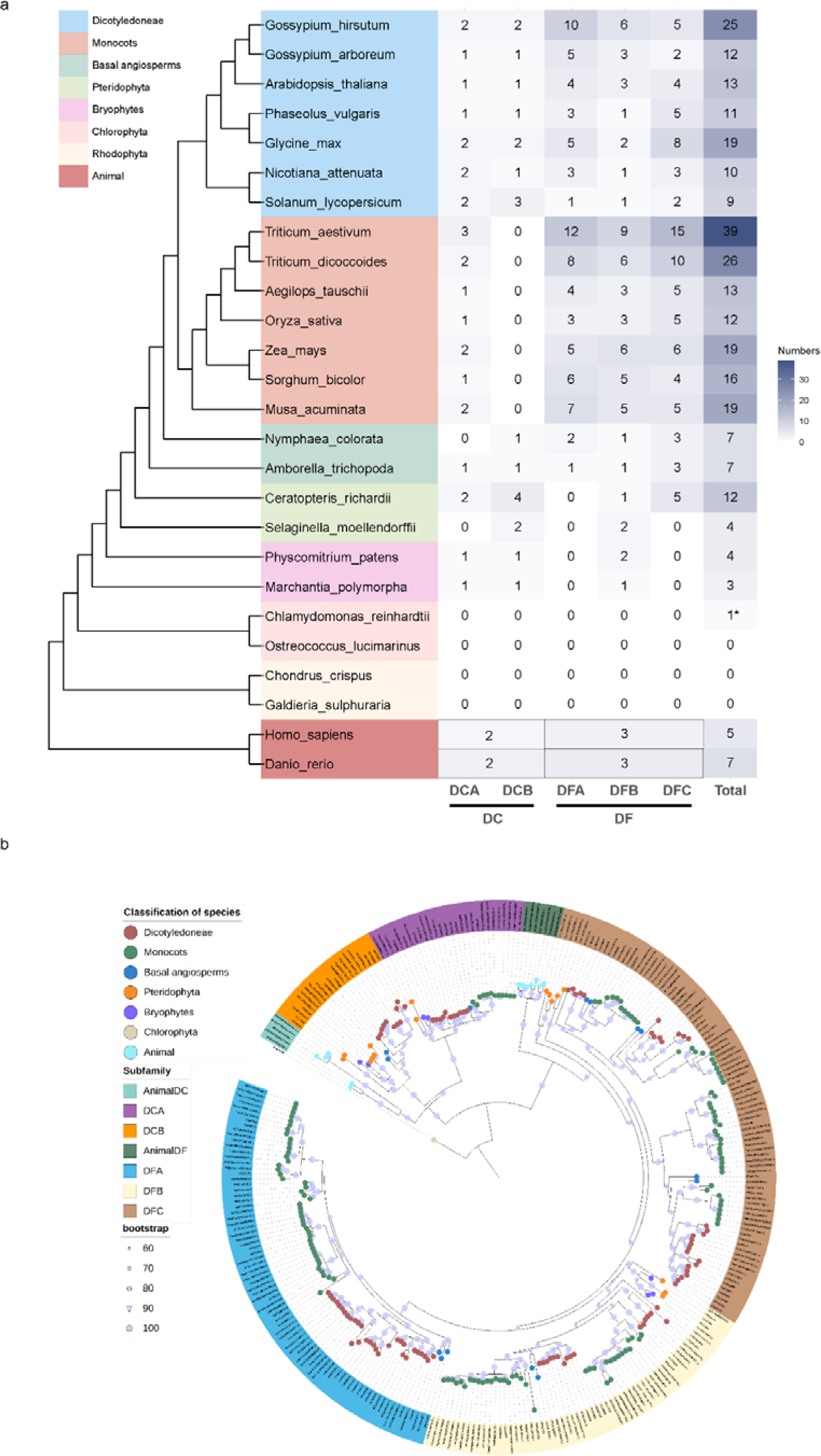
Evolutionary analysis of YTH proteins. (a) Number and classification of YTH proteins identified in 26 species. The evolutionary relationships of these YTH proteins were constructed using the TimeTree (https://timetree.org/). ‘*’ indicates proteins used as outgroups. (b) Evolutionary relationships of YTH proteins. Bootstrap values are represented by circles on the evolutionary branches. The background color of sequence names indicates the classification of YTH subfamilies, and the color of leaf nodes represents the taxonomic units of the species to which the sequences belong.

**Supplementary Fig. 8.**
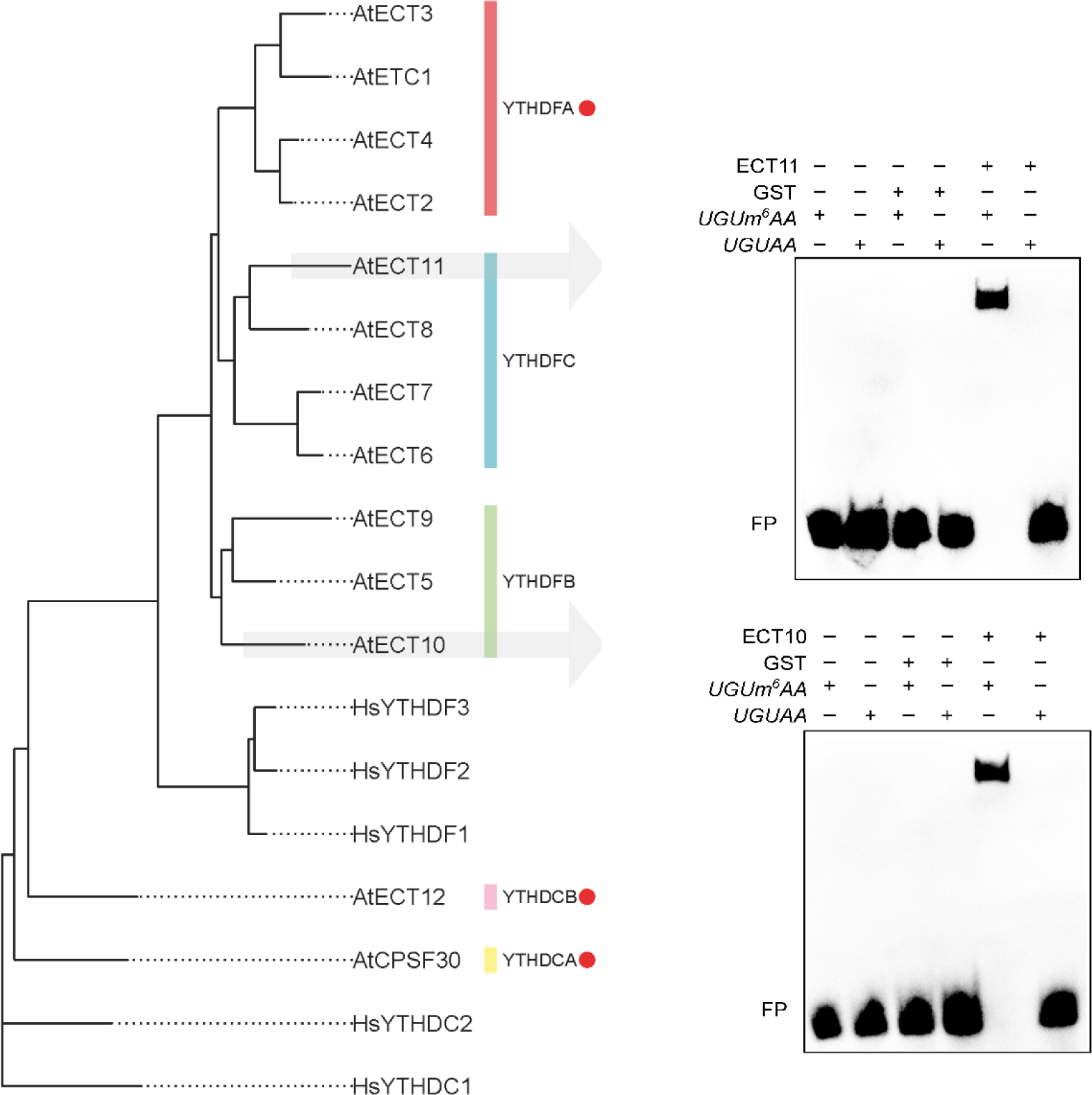
RNA-binding ability of ECT10 and ECT11. **Left**: Phylogenetic tree of YTH proteins in *Arabidopsis* and humans, with subfamilies experimentally confirmed as m^6^A readers highlighted with red circles. **Right**: EMSA assay. 5_-Biotin-labeled synthetic RNA with or without m^6^A modification motif *UGUAA* were incubated with GST-ECT10, GST-ECT11 or GST protein alone (negative control). Samples were analyzed by native PAGE. Gels were blotted onto nylon membranes and signals were detected by streptavidin-coupled horseradish peroxidase and ECL. FP, free probe.

